# Motor training improves coordination and anxiety in symptomatic *Mecp2*-null mice despite impaired functional connectivity within the motor circuit

**DOI:** 10.1101/786822

**Authors:** Yuanlei Yue, Pan Xu, Zhichao Liu, Xiaoqian Sun, Juntao Su, Hongfei Du, Lingling Chen, Ryan T. Ash, Stelios Smirnakis, Rahul Simha, Linda Kusner, Chen Zeng, Hui Lu

## Abstract

Rett Syndrome (RTT) is a severe neurodevelopmental disorder caused by loss of function of the X-linked Methyl-CpG-binding protein 2 (*MECP2*). Several case studies report that gross motor function can be improved in children with RTT through treadmill walking, but whether the MeCP2-deficient motor circuit can support actual motor learning remains unclear. We used two-photon calcium imaging to simultaneously observe layer (L) 2/3 and L5a excitatory neuronal activity in the motor cortex (M1) while mice adapted to changing speeds on a computerized running wheel. Despite circuit hypoactivity and weakened functional connectivity across L2/3 and L5a, the *Mecp2*-null circuit’s firing pattern evolved with improved performance over two weeks. Moreover, trained mice became less anxious and lived 20% longer than untrained mice. Since motor deficits and anxiety are core symptoms of Rett, which is not diagnosed until well after symptom onset, these results underscore the benefit of motor learning.

## Introduction

Experience tells us that we develop skill by repeated efforts at a particular outcome, whether the task is cognitive, physical, or both. Even a relatively simple task such as walking involves a combination of both hard-wired abilities and plasticity for adapting basic movements to different environmental demands while achieving reliable outcomes. The development of motor skills relies heavily on the motor cortex (M1), particularly layers 2/3 and 5 (L2/3 and 5), which undergoes reorganization during learning (*1–5*). Motor learning enhances synaptic connections in M1 by coordinating the clustering of dendritic spines, forming the structural basis for storing motor memory (*6, 7*). L2/3 and 5a provide excitatory input to the major motor output layer, L5b (*8*). The microcircuits in L2/3 integrate sensory feedback, motor planning and primitives into motor output, whereas L5a neurons play crucial roles in action selection, motor control, sequence learning and habit formation (*3, 9*). Neurons in these two layers are reciprocally connected, but we are only beginning to learn how they encode motor skills.

We were particularly interested in understanding how the function of the M1 circuit is altered in a neurodevelopmental disorder known as Rett syndrome (RTT), which is caused by mutations in the X-linked Methyl-CpG-binding protein 2 (*MECP2*) (*10*) and appears to involve a loss of the ability to learn. RTT is striking for the unusual course of the disease: children (typically girls) develop normally until about 18-24 months of age, when they undergo a period of regression wherein they lose acquired motor, cognitive, and social milestones and do not make further gains (*11*). They stop speaking, they develop hand stereotypies that replace purposeful hand use, they develop apraxia and ataxia if they remain ambulatory, and they develop a host of additional neurological symptoms such as seizures, respiratory dysrhythmias, and anxiety. There is no neurodegeneration; in fact, mice in which *Mecp2* is re-expressed in adulthood are completely rescued (*12*). This indicates that there is no fundamental structural abnormality, even though *Mecp2*-deficient neurons in mice are smaller and are more closely packed than in wild-type animals (*13, 14*). MeCP2 expression rises post-natally and seems to have roles in maintaining the physiology of mature neurons (*15–17*), but the *Mecp2-*heterozygous hippocampal circuit was still able to be activated by deep brain stimulation to a sufficient degree to rescue a learning and memory deficit (*18*) and reduce the abnormal hypersynchrony of neuronal activity (*19*).

Considering that the motor cortex also remains structurally intact during regression, we wondered whether the motor circuit was, despite evidence to the contrary, still capable of learning after regression has set in, and specifically whether the motor cortical circuit would respond to a less directed source of stimulation, namely, exercise.

To study M1 activity and particularly cross-layer interaction during motor skill acquisition in wild-type and *Mecp2*-null mice, we monitored L2/3 and L5a of M1 in the mice using two-photon calcium imaging during motor training. The question was what kind of motor learning task would be suitable. Early-symptomatic *Mecp2* null mice are considerably weaker, less coordinated, and less active than wild-type mice (*20*), so voluntary exercise would not work. On the other hand, classic motor learning tasks in rodents, such as operating a lever to get food pellets, depend on operant conditioning, i.e., pairing a specific behavior with a reward. The involvement of reward circuits might therefore complicate experimental investigation of responses in the motor cortex (*21*), as RTT is known to involve abnormalities in mesocorticolimbic reward pathways (*22*). We therefore developed a forced exercise paradigm, where the mice are placed on a computerized running wheel whose speed can be controlled by the experimenter; the skill to be learned is to adapt to changing speeds across multiple training sessions. This experimental paradigm does not require food reward or water restriction, and, since locomotion is partly hard-wired, it requires minimal involvement of the higher-order motor system. While mice underwent two weeks of training on the computerized wheel, we conducted two-photon calcium imaging to record neurons in L2/3 and L5.

We found that training for two weeks reshaped M1 circuit dynamics in conjunction with improvement in motor skill on the wheel. Our evidence supports a model in which motor learning strengthens the connectivity of a subgroup of neurons for storing information while decreasing connectivity among the rest of the population. We also show that despite reduced cross-layer connectivity, a ∼20% overall lower event rate, and impairment in maintaining functional connectivity more than a couple of days, the *Mecp2-null* motor cortex retains sufficient plasticity to support some motor learning. More surprisingly, a mere two weeks of training was able to improve motor coordination on different assays (ladder-walking and accelerating rotarod tests), reduce anxiety (in the open field and elevated plus maze tests), and extend the lifespan of *Mecp2*-null mice by 20%.

## Results

### Male *Mecp2*-null mice are impaired in wheel-running but can improve

We placed symptomatic 7- to 8-week-old, head-fixed, male *Mecp2*-null mice and their wild-type (WT) littermates on a wheel whose speed is controlled by an experimenter (**Fig. 1A**). We analyzed forelimb kinematics while the mice adapted to four different speeds (15, 30, 45, 60 mm/sec). Mic e started at rest (which we will call “speed 0”), th en the speed was increased by 15 mm/sec every two minutes up to 60 mm/sec, followed by a two-minute rest period, then the speed was set to 60 mm/sec and decreased by 15 mm/sec every two minutes down to 0 mm/sec (**Fig. 1A**, right panel). This experiment was performed daily over 12-14 consecutive days. Since the limbs of head-fixed mice do not support the full weight of the body, the range of motion of the forelimbs was quite large. (This could make the task more challenging, but by relieving the mice of the need to support their full body weight, it might also help the null mice, who can suffer exercise fatigue (*23*).) We used DeepLabCut (*24*) to train a deep neural network to track several parameters for each step cycle (**Fig. 1B and fig. S1A**). The movement of each paw is automatically tracked by the algorithm. Step number is reported for the left forepaw, contralateral to the site of imaging in area M1 on the right, but both front paws showed similar movements. The most reliably detected features of each step were the rising phase of a spike in the horizontal plane (x) and a complete spike in y (the vertical plane), and these were used in the analysis to represent each step (**Fig. 1C**).

**Fig. 1.**
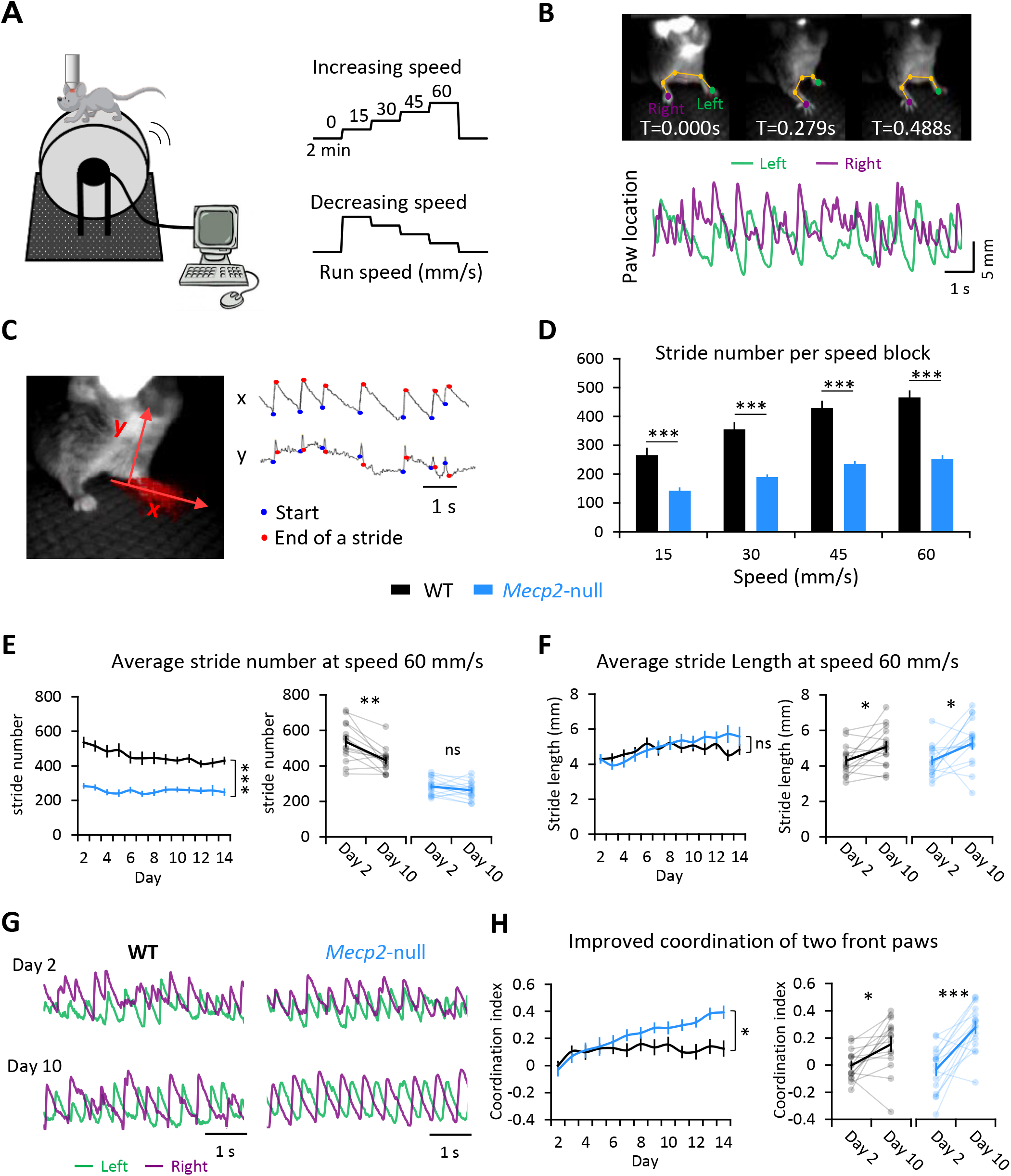
Following the development of motor skill in forced wheel running. (**A**) The experimental set-up. A mouse, whose head is fixed under a two-photon microscope, runs on the computerized wheel, whose speed is controlled by the experimenter. (**B**) *Top*: Video frames of forepaw movements at three representative time points, with paw locations labeled by DeepLabCut. *Bottom*: Sample temporal traces for the left and right paws of one mouse moving at 60 mm/s on the wheel. The paw location at any given moment is calculated as the distance between the median paw location and location at that time. (**C**) Automated stride analysis. X and Y axes represent horizontal and vertical paw movements, respectively. A step is represented by the rising phase of a spike in X and a complete spike in Y. (**D**) The average number of strides completed by the left paw at each speed (during increasing-speed mode) over the training period. Error bars represent mean ± SE. n=13 mice for WT group; N=15 for *Mecp2*-null group. *** *P* <0.001, RM-ANOVA with Sidak’s Post-Hoc. (**E**) Stride number (left) and stride length (right) at speed 60 mm/sec over 14 successive days. (**F**) *Left.* Comparison of stride length between genotypes every 10 seconds during the 2-min block of speed 60 mm/s on days 2 and 10, fitted with a straight line. *Right:* within-genotype comparisons of stride length during early learning (day 2) and consolidation (day 10) phases. (**G**) Sample temporal traces for the forepaws of a mouse running at speed 60 mm/s shows improved coordination (anticorrelation) by day 10. (**H**) Coordination at speed 60 mm/s improves over the 14-day training period. The coordination index was calculated as the negative value of the correlation coefficient of the two forepaw traces during the 2-min block of speed 60 mm/s. N=13 mice for WT group; n=15 for *Mecp2*-null group. * *P* <0.05, RM-ANOVA test.

The *Mecp2*-null mice had great difficulty adapting to the moving wheel: their limbs repeatedly “froze” (**fig. S1B,** indicated by the red arrows), possibly due to either muscle weakness (*23*), the apraxia that is seen in RTT, or anxiety from being on an unpredictable wheel. Because of this, *Mecp2-null* mice took fewer strides than their WT counterparts at all speeds tested, but the magnitude of the change in stride number from lower to higher speeds was very similar between the two genotypes (an increase of approximately 70% from 15 to 60 mm/sec; WT, 15 mm/sec: 271.8±25.1; WT, 60 mm/sec: 475.7±23.7; Null, 15 mm/sec: 146.0±9.5; Null, 60 mm/sec: 261.5±12.5. **Fig. 1D**). As the WT mice developed skill on the wheel, they took fewer (538.8±29.6 on Day 2 vs. 434.1±20.9 on Day 10) but slightly longer strides (4.1±0.2 mm on Day 2 vs. 5.0±0.3 mm on Day 10)(**Fig. 1E-F**). This pattern suggests that, in WT mice, the process of learning to adapt to the running wheel takes about one week (the early learning phase), and that the learned skill is consolidated in the second week (consolidation phase). The null mice did not show a significant reduction in stride number (perhaps because moments of “freezing” skewed the average), but their average stride length increased steadily over the course of training (**Fig. 1E-F**). The most striking change, however, was that the *Mecp2*-null mice developed much better forelimb coordination, reflected by the increased negative correlation of the traces of the two front paws (**Fig. 1G and H**, **videos 1 and 2**). Like the wild-type mice, the null mice showed greater changes during the first week of training than the second week (**fig. S1C**), suggesting they follow a similar pattern of learning and consolidation, though they continued to improve during the second week (**Fig. 1H**). It may be that loss of MeCP2 extends the period required for motor skill learning but does not abolish it. We next investigated how the motor circuit might be reshaped by this training.

### M1 neurons in *Mecp2*-null mice are hypoactive but retain plasticity

Although *ex vivo* experiments show that cortical pyramidal neurons lacking *Mecp2* are hypoactive (*25*), we wondered how M1 pyramidal neurons behave *in vivo* during movement. In particular, we wanted to know how the interaction between the pyramidal neurons in layers 2/3 and 5a is refined by motor training. Previous studies have been limited by the inability to record neurons in the two layers simultaneously, but here we used a specialized two-photon microscope equipped with high-speed piezo-Z optical-axis objective positioner that was able to accomplish this feat **(Fig. 2A**). We confirmed the proper depth for L5a with the L5-specific Rbp4-Cre line (**fig. S2A**, top) and did not observe obvious depth differences between the WT and *Mecp2*-null mice (**fig. S2A**, bottom). Thus, although the greater depth of L5a diminishes the quality of recording, it does so to the same degree in both genotypes and so did not affect comparisons between the two mouse lines. L5a somata were significantly larger than those of L2/3 (**fig. S2B**), confirming the different types of neurons imaged in these two layers.

**Fig. 2.**
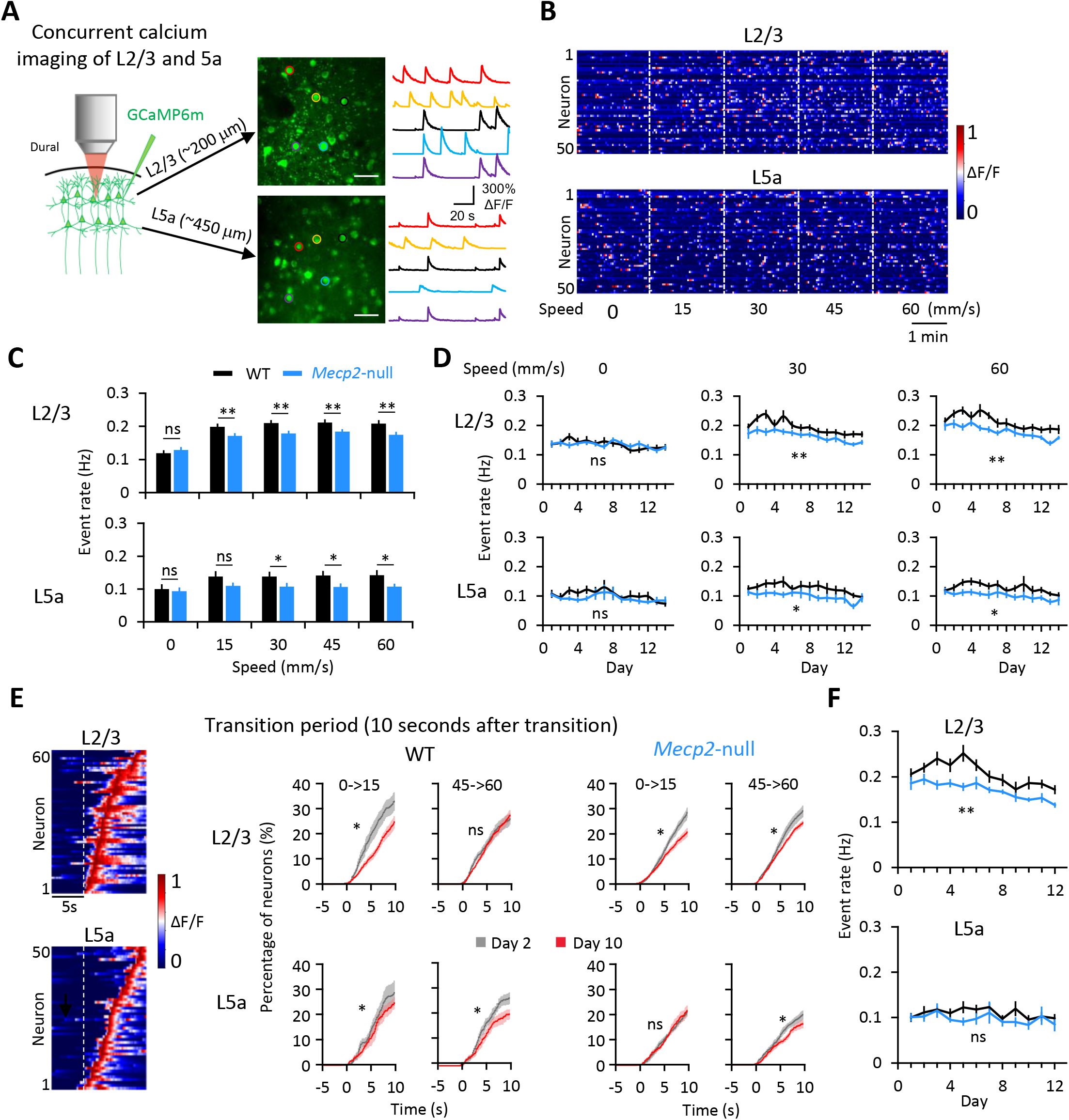
Motor learning streamlines distribution of event rates in L2/3 and 5a. (**A**) Images of pyramidal neurons expressing GCaMP6m in L2/3 (top) and L5a (bottom) of one mouse. The color of the circle around a given neuron in the photo matches the color of its activity trace. (**B**) Heatmap of neuronal activity from 50 representative neurons (rows) during increasing- speed mode. Colors represent normalization to the highest fluorescent intensity change (ΔF/F) of each neuron. (**C**) Averaged event rates of the detected neurons at five speeds. (**D**) Averaged event rates across training days. L2/3 WT: n=13; L2/3 *Mecp2*-null: n=14; L5a WT: n=11; L5a *Mecp2*- null: n=13. * *P* < 0.05, ** *P* < 0.01, RM-ANOVA test. (**E**) *Left*, Heatmap of neuronal activity around a speed transition from 0 to 15 mm/sec from one WT mouse. *Right*, Cumulative proportion of transition-associated neurons from all mice, on days 2 and 10, sorted by the peak of their response during the 10 seconds after the transition from 0 to 15 and 45-to 60 mm/s. * *P* <0.0001, Kolmogorov- Smirnov test. (**F**) Averaged event rate of neurons during 10 seconds after five speed-increasing transitions on each day of training. L2/3 WT: n=11; L2/3 *Mecp2*-null: n=12; L5a WT: n=8; L5a *Mecp2*-null: n=10. RM-ANOVA test. ns, not significant. Error bars represent mean ± SE.

We identified the fluorescent transients (events) during the task (**fig. S2C, D**), and analyzed the event rate of the same groups of pyramidal neurons in the right forelimb area of the primary motor cortex (M1) across the 14 days of imaging. In wild-type mice, the mean event rate in both L2/3 and L5a was higher during running than at rest (**Fig. 2B-D and fig. S3**). Null mice showed similar L2/3 and 5a event rates as WT at rest and 15 mm/s, but both layers were less active at all other speeds (**Fig. 2C, D**). The differences did not depend on the direction of change (i.e., increasing or decreasing from the previous block, **figs. S3, S4**). The event rate of L2/3 neurons fell during running by about 15-20% after learning in L2/3 (from 0.23 ± 0.01 on Day 1-2 to 0.20 ± 0.01 on Day 9-10 for WT, p = 0.010; and from 0.21 ± 0.01 on Day1-2 to 0.17 ± 0.01 on Day 9- 10 for the null group, p = 0.023, Wilcoxon signed-rank test), but not in L5a (from 0.11 ± 0.01 on Day 1-2 to 0.12 ± 0.01 on Day 9-10 for WT, p = 0.779; and from 0.11 ± 0.01 on Day1-2 to 0.10 ± 0.01 on Day 9-10 for the null group, p =0.241, Wilcoxon signed-rank test test) (**fig. S5A**). The overall distribution of event rates shifted toward lower rates after the learning phase, and the peak narrowed at both rest and running states (**fig. S5B, C**), reflecting less variation in stimulus coding. The evolution of event rates corresponded with the mouse’s development of skill on the running wheel (**Fig. 1**), consistent with previous findings demonstrating less cortical activation when executing a well-practiced as opposed to new motor behavior (*26–29*).

We next examined whether the M1 circuit in wild-type or null mice responded to changes in wheel speed. We found that a portion of the L2/3 and L5a ensembles responded with a higher event rate in the 10-second period after the change than in the 5-second period prior to the change (**Fig. 2E**). Comparing these transition-associated responses at each transition between day 2 (learning phase) and day 10 (consolidation phase), we found that after learning, ∼20% fewer L2/3 neurons in both WT and *Mecp2*-null mice responded to the initial transition from rest (speed 0) to 15 mm/s, and fewer L5a neurons (∼25% for WT and ∼15% for *Mecp2*-null) responded to transitions between running speeds (**Fig. 2E**). Overall, the difference was greatest from 0 to 15 in L2/3 for both animals, but the evolution of learning-induced changes in the response to speed changes was similar between WT and *Mecp2*-null mice (**Fig. 2E**). The event rates of *Mecp2*-null L2/3 neurons, but not L5a, were significantly lower than WT during the10- second period after a speed change (**Fig. 2F**). When we identified specific neurons and followed them over multiple days (**fig. S6A**; see Methods) to examine how they respond to speed changes, we found that they were exceptionally plastic: a given neuron might respond (or not) to a transition on any given day (**fig. S6B, C**). This plasticity of response to a transition was at the same level in the WT and *Mecp2*-null mice (**fig. S6D**). The response to speed changes thus takes place at the level of the ensemble, not the individual neuron.

### Motor learning strengthens functional connectivity for a small proportion of neuronal pairs

Information is encoded at the circuit level through the coordinated firing patterns of neurons in the ensemble (*30*). We therefore analyzed the functional connectivity between recorded neurons in the M1 network throughout training using correlation analysis (*31*). Functional connectivity is commonly estimated by Pearson correlation coefficients, which include direct as well as indirect correlations. Direct correlation reflects functional connectivity within the local circuit by excluding shared input from afferent brain regions (*8*).

We first calculated the Pearson correlation coefficient between each pair of neurons within L2/3 and L5a, as well as across these two layers in both WT and *Mecp2*-null mice. At rest, the correlation coefficients of neuronal pairs did not differ between the learning and consolidation phase, with the exception of layer L2/3 neurons in the null mice (**fig. S7A**). There were no differences in correlations either within or between layers at any running wheel speeds from learning (days 1-2) to consolidation (days 9-10) (**fig. S7B**). However, in both genotypes, the L2/3 neuronal pairs were less functionally connected during the consolidation phase. The overall functional connectivity of neuronal pairs during running, both within L2/3 and across the two layers, diminished with training (**Fig. 3A**, top row). The distribution of correlation coefficients was rather broad (between -0.5 and 1) across the population of M1 neurons, and shifts in the distribution of pairwise correlations can be seen across training (**fig. S8**). Yet in the 5% of neuronal pairs that were most strongly connected—i.e., those in which the Pearson correlation coefficient was larger than the mean plus two times the standard deviation of the correlation coefficient—functional connectivity actually increased with training (**Fig. 3A**, bottom row, and **fig. S8)**. These results suggest that refining a motor skill (in this case, adapting one’s gait to different speeds of a surface controlled by someone else) increases functional connectivity between already-correlated neurons, with an overall net decrease in functional connectivity across the rest of the population, within L2/3 and across L2/3 and 5a. This circuit refinement occurred in the *Mecp2*-null mice as well.

**Fig. 3.**
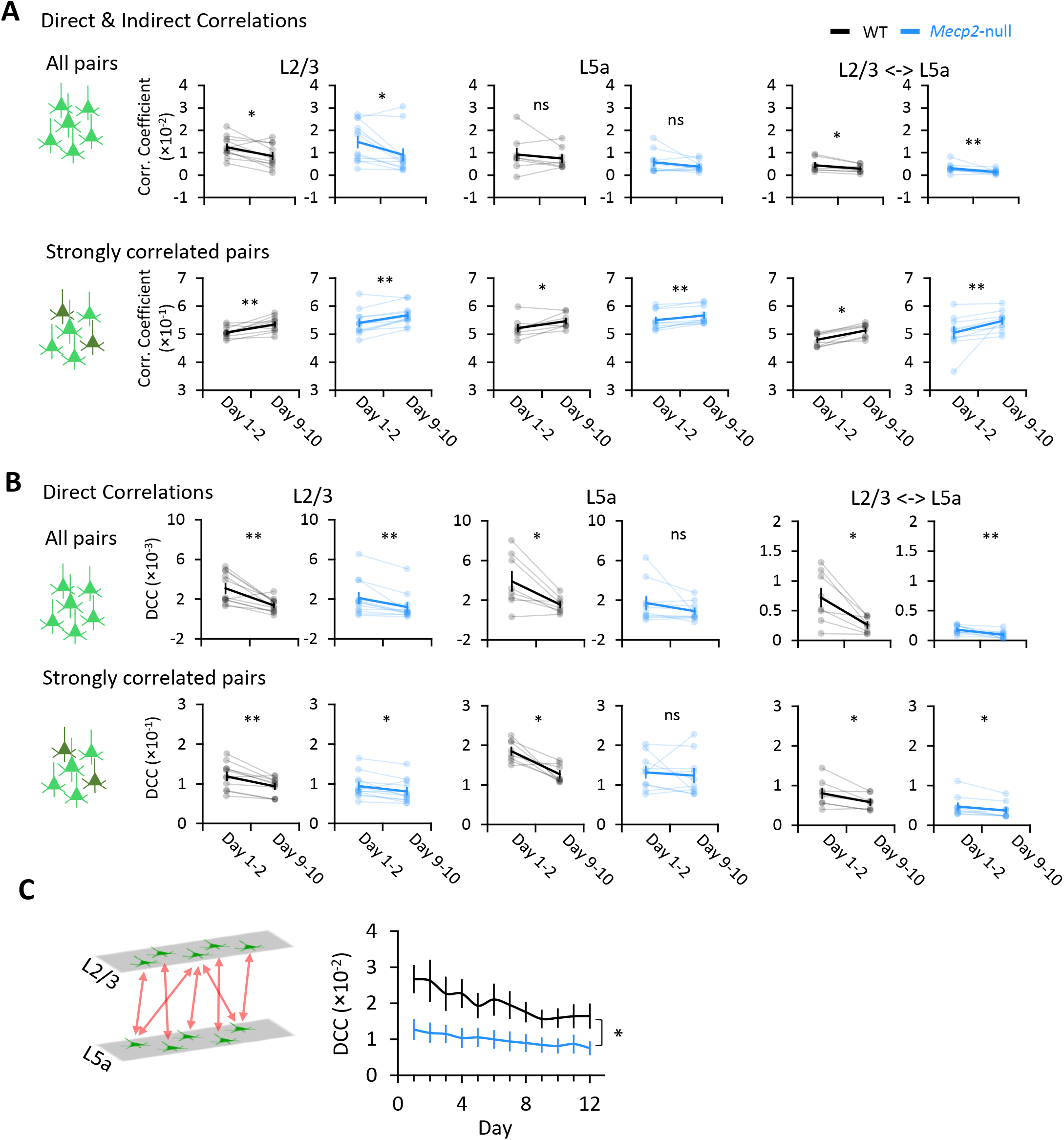
Overall functional connectivity across M1 decreases with learning, but a small percentage of neuronal pairs become more strongly correlated. (**A** and **B**) Mean direct and indirect correlation coefficients calculated as Pearson correlation coefficients (**A**) and direct correlation coefficient calculated using DCA (Direct Correlation Analysis) (**B**) between pairs of neurons within L2/3, within L5a, and across both layers during four running blocks for an individual WT or *Mecp2*-null mouse (top: all pairs of correlation; Bottom: strong correlations, defined by a correlation coefficient that was greater than twice the standard deviation). Light lines connect data from each individual mouse; dark lines connect the averaged data from all mice. L2/3 WT, n=11 mice; L2/3 *Mecp2*-null, n=12 mice; L5a WT, n=8 mice; L5a *Mecp2*-null, n=10 mice (we excluded the L5a data from mice from which the number of neurons detected in L5a was less than 20). Across both layers, WT, n=8 mice; *Mecp2*-null, n=10 mice. Error bars represent mean ± SE. ns, not significant; * *P* < 0.05; ** *P* < 0.01; *** *P* < 0.001,Wilcoxon signed-rank test. (**C**) Direct correlation coefficients (DCC) of neurons across M1 layers 2/3 and 5a. WT, n=8 mice; *Mecp2*-null, n=10 mice. Error bars represent mean ± SE. * *P* < 0.05, RM- ANOVA test.

To verify that the changes we detected in the M1 circuit bore a causal relationship with motor training, we also imaged age-matched training-naïve mice during five minutes of free-running (i.e., on a frictionless wheel), on two separate days a week apart. There was no change in the correlation coefficients of any neuronal pairs in mice of either genotype between days 1 and 8 (**fig. S9A**). We then imaged trained mice of both genotypes during a free-run mode before training on days 1-2 and 9-10. The correlation coefficients dropped significantly for L2/3 neurons in both WT and *Mecp2*-null mice, and across both layers in *Mecp2*-null mice (**fig. S9B**). These data demonstrate that the changes observed in M1 result from motor learning, not merely from the activity of running or from the passage of time.

To determine whether the heightened functional connectivity of certain neuronal pairs might be due to shared input from afferent brain regions, we employed a maximum entropy-based inference method (Direct Correlation Analysis, DCA) that has been used to infer direct interactions from biological datasets such as gene expression data or sequence ensembles while excluding indirect interactions (*32–34*). We calculated the averaged direct correlation coefficient (DCC) and found that the DCC decreased with training for all pairs, including strongly correlated pairs, within L2/3, within L5a, and across these two layers (**Fig. 3B**, top row). The magnitude of the decrease was slightly smaller in the *Mecp2*-null mice than in WT mice. This indicates that the functional connectivity among L2/3 neurons decreased over the course of learning. The data also indicate that the greater temporal correlation of firing among a small proportion of L2/3 neurons (**Fig. 3B**, bottom row) was due to more synchronized input to these neurons from other brain regions (most likely the somatosensory and premotor cortex) over the course of learning (*8*). It’s worth noting that, after learning, there was no difference between genotypes in direct correlation (**fig. S10**), but cross-layer connectivity between L2/3 and L5a was much lower in the *Mecp2* null mice than in the WT mice (**Fig. 3C**), reflecting a profound disruption of interlaminar functional connectivity in these animals.

In sum, functional correlations between strongly correlated neuronal pairs increase with learning, likely due to synchronized input from the somatosensory and premotor cortices. This small portion of strengthened functional correlations takes place against a background of overall diminishing functional correlations within the ensemble. This suggests that the ensemble becomes more efficient by recruiting the same small group of neurons to the task.

### MeCP2-deficient neuronal pairs are less stable than WT and do not show proximity bias

To further explore the effect of MeCP2 deficiency on M1 circuit plasticity, we used DCA to search for functional connectivity among the same group of neurons within each mouse. We were able to track ∼20% of all active neurons throughout the 12 days using the method in **fig S6A** and counted how many times we detected functional connectivity between the same pair of neurons (**Fig. 4A**). Strongly correlated pairs of neurons were more likely to be detected repeatedly over the two-week period in the wild-type mice than in *Mecp2*-null mice (**Fig. 4, B and C**); the persistence of a given pairing was more probable with running than rest in wild-type mice (**Fig. 4C**). Strikingly, the majority of pairs were detected in WT mice over four days (during running), but in *Mecp2*-null mice, most neuronal pairings occurred only once or twice (during running or resting). Functional connections in both genotypes were virtually undetectable beyond 8 days. The number of pairs that formed in wild-type mice on only one or two days during runs was half the number found in the null; conversely, null mice had about half the number of pairs as WT mice (during rest or running) that were detected more than four days (**Fig. 4C**, right panel). In other words, the null mice are able to form functional connections but at half the rate and for not nearly as long. Although we were not examining actual synaptic connections, these data are in agreement with previous studies showing that physical synaptic connections are attenuated in *Mecp2*-null mice (*35, 36*).

**Fig. 4.**
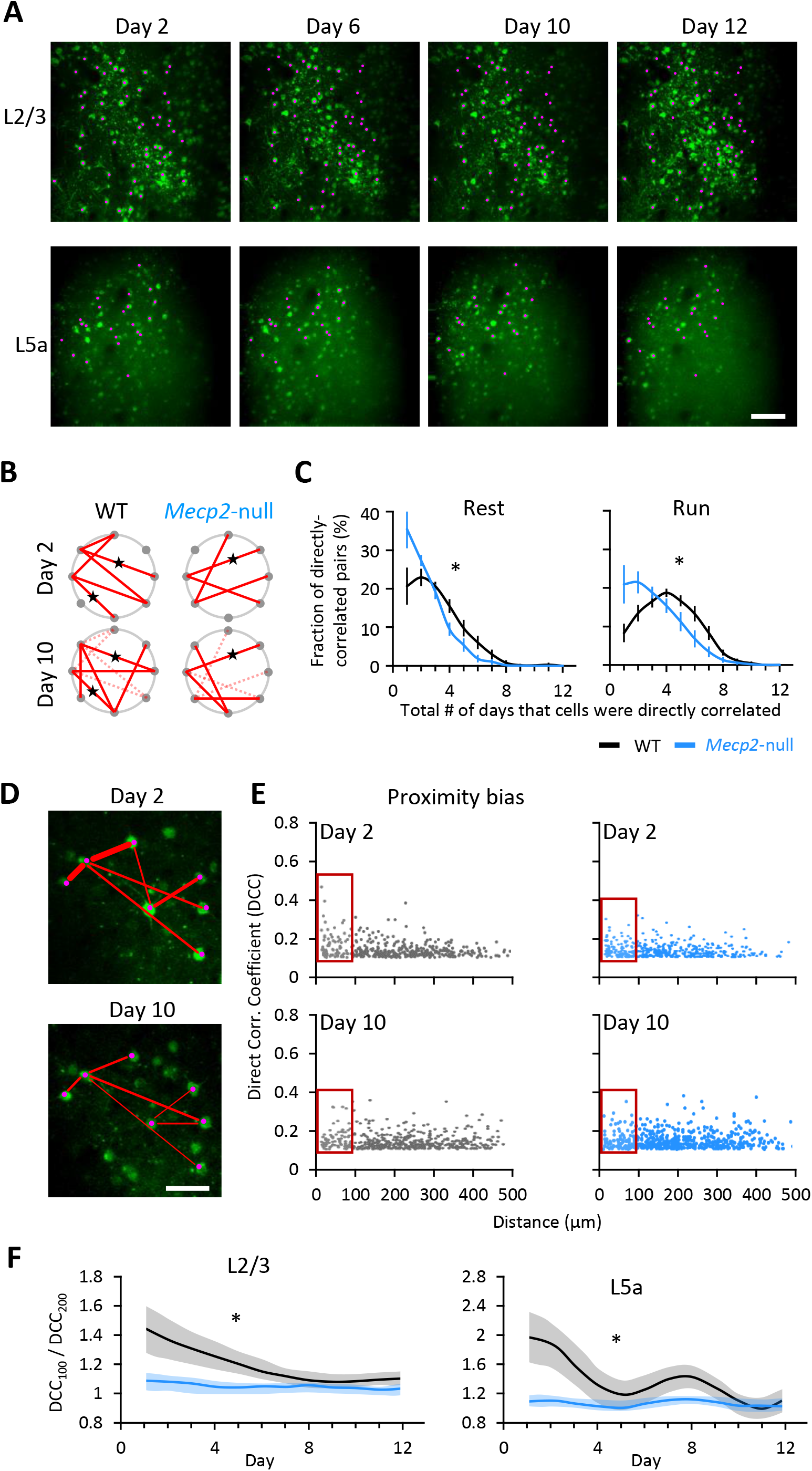
Loss of *Mecp2* shortens the duration of synaptic connections during learning. (**A**) Images of same field of view on L2/3 and 5a on different days from a WT mouse. Pink dots indicate the same active neurons detected on multiple days. Scale: 100 μm. (**B**) Diagram demonstrating direct correlation between neurons on day 2 (top) and day 10 (bottom) of WT (left) and null mouse (right). Each dot represents one neuron. Red lines indicate direct correlations between two neurons. Dashed lines indicate connections that disappeared from day 2 to day 10. Stars label the connections that existed on both days (there were fewer such connections in Null mice than in WT mice). (**C**) The percentage of the directly correlated neuronal pairs that were detected over a different number of days across the two-week testing period at rest (left) and running (right) state. WT, n=9 mice; *Mecp2*-null, n=9 mice. Error bars represent mean ± SE. * *P* < 0.05, t-test between the kurtosis values that represent the shape of two curves. (**D**) Direct correlations between neurons in the same field of view on days 2 and 10. Red lines represent direct connectivity between a pair of neurons. Line thickness represents correlation strength. Scale: 50 μm. (**E**) Relationship plot between the direct correlation coefficient (DCC) of a pair of neurons on days 2and 10 and their physical distance from each other. X and Y axes represent the distance of the paired neurons and their DCC, respectively. The red box marks the neuronal pairs whose distance is less than 100 µm. (**F**) Ratio of DCC between neuronal pairs with a distance less than 100 µm (DCC100) and between 100-200 µm (DCC200). L2/3 WT, n=11 mice; L2/3 *Mecp2*-null, n=12 mice; L5a WT, n=8 mice; L5a *Mecp2*-null, n=10 mice, * *P* < 0.05, RM- ANOVA test.

We next asked whether functional connectivity was influenced by physical proximity, as previous studies have observed a ’proximity bias’ in normal circuits (*37, 38*). In the initial stages of motor training, we found that the closer that two L2/3 neurons were to each other in the WT brain, the higher their direct correlation coefficient (**Fig. 4, D and E**). This proximity bias was much less apparent in the null mice, similar to what we previously observed in the null hippocampal circuit (*19*) **(Fig. 4E**). To visualize how proximity bias evolved with training, we plotted the ratio of the direct correlation coefficient for pairs closer than 100 μm (DCC100) to pairs at a distance between 100 μm and 200 μm (DCC200) (**Fig. 4F**). In WT mice, this ratio declined over the first week of training, then stabilized in L2/3 and L5 (**Fig. 4F**, black lines). In *Mecp2*-null mice, however, the ratio started low and did not change with training (blue lines). Thus, WT mice have a proximity bias that diminishes with learning, as the strongly correlated pairs whose functional connectivity grows are not necessarily near one another. *Mecp2-*null animals have no proximity bias to begin with, however, despite the fact that they have smaller, more closely packed neurons (*13, 14*).

### Trained *Mecp2* null mice are better coordinated, less anxious, and live longer

Daily running on a speed-controlled wheel is analogous to forced exercise, which has proven broadly beneficial in Parkinson’s disease patients (*39, 40*). Exercise has also been shown to reduce stereotypic behaviors in children with autism spectrum disorders (ASD) (*41*). We therefore asked whether the variable-speed wheel training improved motor function more generally in *Mecp2*-null mice. We first examined the performance of the mice crossing a ladder over 20 trials after the last day of training on the wheel (**Fig. 5A**). As expected, age-matched naive *Mecp2*-null mice had difficulty throughout the 20 trials, though they did improve. The trained *Mecp2*-null mice, however, performed like WT after about 12 trials of ladder running (**Fig. 5B and fig. S11**). Compared to the naïve *Mecp2*-null mice (**video 3**), trained *Mecp2*-null mice made significantly more correct steps (steps between adjacent high rungs) and fewer irregular steps (i.e., those not between high rungs; see Methods) (**Fig. 5C, video 4**). Trained *Mecp2*-null mice maintained their superior coordination 2 days and 4 days after the last day of wheel running, indicating that benefits from assisted motor training can last at least a few days (**Fig. 5D**).

**Fig. 5.**
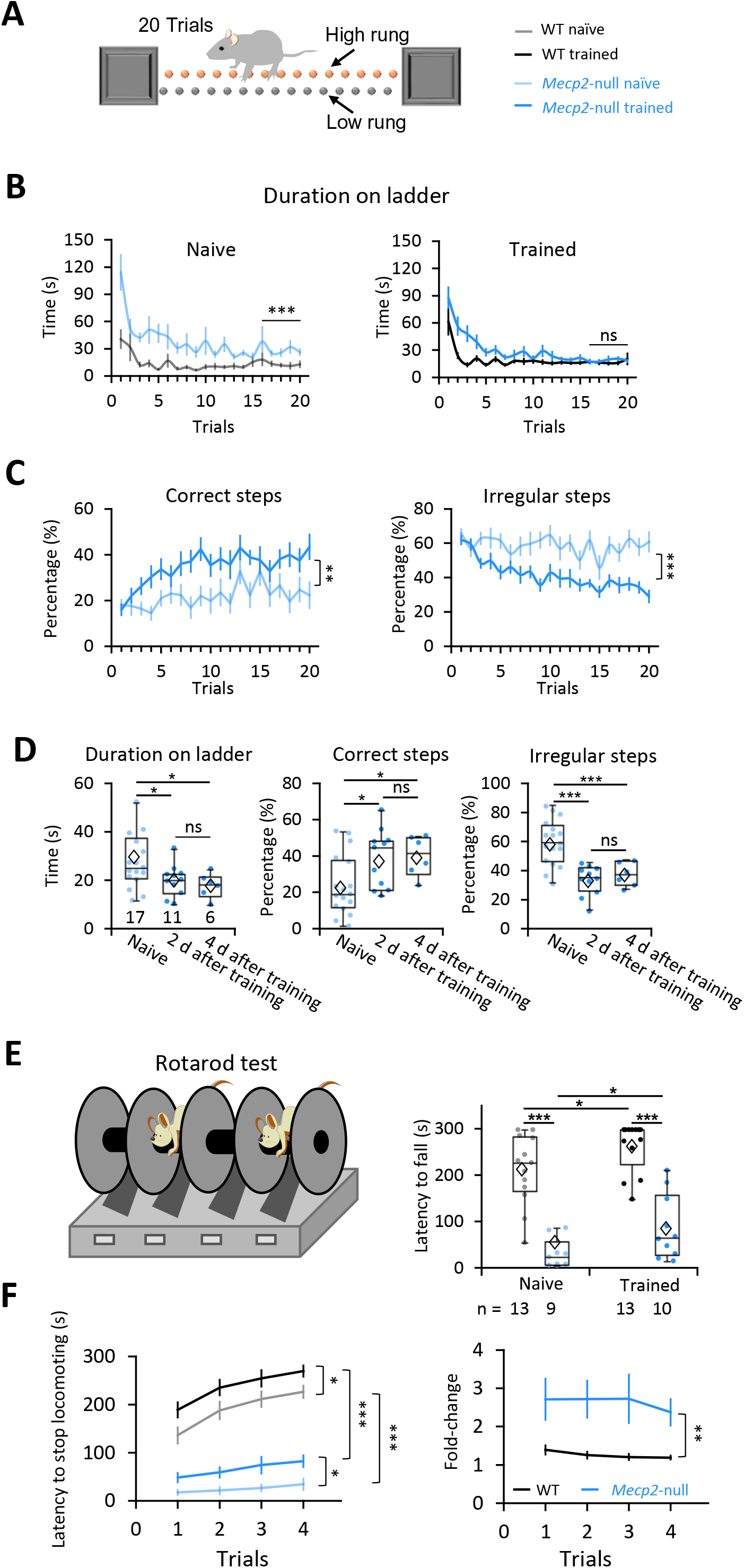
Motor learning improved coordination of *Mecp2*-null mice in different assays. (**A**) The ladder test as a measure of coordination. The animals run from one end to the other 20 times (20 trials). (**B**) How much time it took mice to walk from one end of the ladder to the other. Naive: WT: n=15; *Mecp2*- null, n=17; Trained: WT: n=19; *Mecp2*-null, n=17. (**C**) Averaged percentage of correct and irregular steps at each trial. In B and C: ns, not significant, ** *P* < 0.01, *** *P* < 0.001, RM-ANOVA test. (**D**) Performance parameters from the last five trials of the ladder test 2 or 4 days after wheel-training ended (for trained mice) or from age-matched naïve mice. Naive: n=17 mice; 2 day after training: n=11 mice; 4 day after training: n=6 mice. Error bars represent mean (diamond) ± SE. * *P* < 0.05, *** *P* < 0.001, One-way ANOVA test with Fisher’s LSD post hoc. (**E**) Latency to fall in the first trial of the accelerating rotarod test 2 days after training. The number of mice examined is presented in each column. Error bars represent mean ± SE. ns, not significant, *P* < 0.05, *** *P* < 0.001, Two-way ANOVA test with Sidak’s Post-Hoc. (**F**) Latency to stop locomotion (left) and fold-change (right, calculated as the mouse’s latency to stop divided by the mean value of the naïve mice) on the rotarod for 4 trials. Naive: WT: n=11 mice; *Mecp2*-null, n=7 mice; Trained: WT: n=9 mice; *Mecp2*-null, n=8 mice. Error bars represent mean ± SE. * *P* < 0.05, ** *P* < 0.01, *** *P* < 0.001, RM-ANOVA test.

To see whether this improved coordination translated to other circumstances, we also tested mice on the accelerating rotarod test. (Although this test is similar to the computerized running wheel, the mice bear their own full weight.) Motor training more than doubled the latency to fall in *Mecp2*-null mice, though they still performed much more poorly than their trained WT littermates (**Fig. 5E**). We noticed that, instead of falling off the rod, most mice simply stopped running and held onto the rotating rod when they got tired, perhaps reflecting the fatigue to which *Mecp2*-deficient muscles are vulnerable (*23*). We therefore decided to compare the time when the animals first stopped locomotion, either by holding onto or falling off the rod. Although motor training significantly increased the latency to stop locomotion by either means, for trained mice of either genotype compared to naïve controls, (**Fig. 5F, left**), the fold-change was much more notable in the *Mecp2*-null mice than that in WT mice (**Fig. 5F, right**).

The trained mice did not show greater strength in the grip test, so their improved motor function was not due to an increase in muscle strength (**fig. S12A**). Nor did they move faster or longer distances in the open field test (**fig. S12B**), but the trained *Mecp2*-null mice spent significantly more time in the anxiogenic center area than untrained *Mecp2*-null mice (**Fig. 6A**). This suggests that forced motor training mitigates the *Mecp2*-null mouse’s well-documented anxiety (*15, 42*). In another classic assay for anxiety, the elevated plus maze test, the naïve null mice spent more time in the open arms than WT even though they did not enter them as frequently (**Fig. 6B**, left), as others have observed (*42*). Trained *Mecp2*-null mice entered the open arms more frequently than the closed arms, signifying less anxiety (**Fig. 6B, right**).

**Fig. 6.**
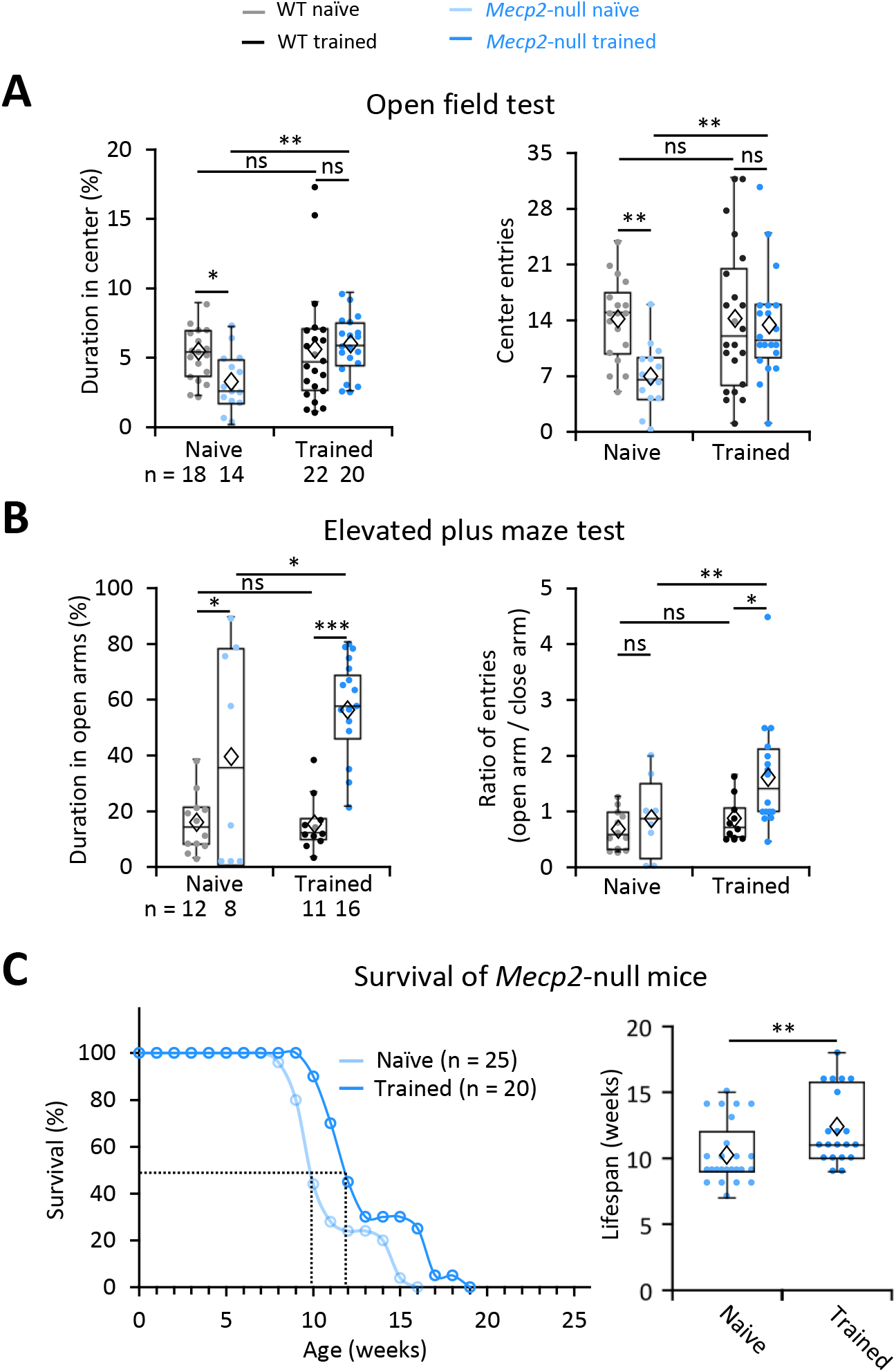
Motor training diminished anxiety and extended survival of *Mecp2*-null mice. (**A**) Performance in the open field test. Naive: WT: n=18 mice; *Mecp2*-null, n=14 mice; Trained: WT: n=22 mice; *Mecp2*-null, n=20 mice. Error bars represent mean ± SE. ns, not significant, * *P*< 0.05, ** *P* < 0.01, *** *P* < 0.001, Two-way ANOVA test with Sidak’s Post-Hoc. (**B**) Performance in the elevated plus maze test. Naive: WT: n=12 mice; *Mecp2*-null, n=8 mice; Trained: WT: n=11 mice; *Mecp2*-null, n=16 mice. Error bars represent mean ± SE. ns, not significant, * *P* < 0.05, ** *P* < 0.01, *** *P* < 0.001, Two-way ANOVA with Sidak’s Post-Hoc. (**C**) *Left,* Exercise training extended the survival of *Mecp2*-null mice. *Right*, Average lifespan of naïve and trained *Mecp2*-null mice in the plot at left. Naive: n=25 mice; Trained: n=20 mice; ** *P* < 0.01, Mann-Whitney U test. In the box plots, diamonds represent the group mean.

Lastly, the trained mutant mice survived ∼20% longer (from 10 to 12 weeks; Naïve: 10.2 ± 0.5 weeks vs. Assisted WR: 12.3 ± 0.6 weeks; *p* = 0.002, Mann-Whitney U test; **Fig. 6C**). We conclude that forced motor training can improve coordination, rescue anxiety, and extend the survival of *Mecp2*-null mice. When we analyzed the relationship between survival time and improved performance on different assays, we found no correlation between lifespan and better performance on the rotarod or anxiety tests, but we did find a weak correlation (p = 0.236 and r = 0.336; **fig. S13**) between coordination and lifespan. This suggests that motor learning may contribute to extending the lifespan of *Mecp2*-null mice beyond the benefits of exercise itself. This would align with previous studies indicating that environmental enrichment benefits Rett mice (*43–45*).

## Discussion

The idea that practice enables the refinement of motor skills is uncontroversial, but the motor circuit changes that underlie this intuitive principle have been challenging to elucidate. Our results support a model in which motor skills are learned by judiciously strengthening the functional connectivity of a subgroup of neurons in order to store information while decreasing connectivity among the rest of the population to maintain flexibility for learning new skills (*46*). This reorganization is attenuated but not abolished by loss of MeCP2. The motor circuit in RTT, although hypoactive and less able to maintain synaptic connections, retains sufficient plasticity to support learning, even though the entire brain is disturbed by loss of MeCP2 in these animals.

It has been proposed that individual neurons increase their ability to discriminate similar stimuli through three processes during motor learning: sharpening their tuning curves, gain modulations, and peak shift (*47, 48*). Previous studies on primate motor and sensory cortices revealed that motor learning is restricted to a subgroup of neurons (*49, 50*). Early training modulates the activity of both the target neurons and nearby cells; as training progresses, only the activity of the target neurons is modulated. Our observation that motor training strengthened the functional connectivity of a small proportion of neurons is consistent with these studies and supports the recently proposed notion that motor learning leads to refinement of cortical population dynamics so that more reliable neural trajectories produce skilled behavior (*51*). Interestingly, however, only the indirect, not direct, correlation coefficients of strongly correlated pairs increased. We hypothesize that this change in functional connectivity is due to a specific increase in the strength of the input from the somatosensory cortex (S1) to L2/3. The main inputs to L2/3 of M1 are from the somatosensory cortex and premotor cortex (*52*), but motor learning in the present study depends to a much greater degree on somatosensory feedback coming from S1 rather than volitional inputs from the premotor areas. Moreover, the L2/3 M1 neurons do not respond specifically to running speeds or speed transitions, suggesting that their activity may be more related to somatosensory feedback and other task-related signals. Further investigation is needed to determine whether afferent axons from S1 form more connections with the highly connected M1 neurons after learning.

A recent study that recorded L2/3 neurons in mice grasping for food on a rotating wheel concluded that these neurons respond to success or failure in the task, rather than specific elements of the movements themselves, in order to reinforce skill learning (*53*). It may be that the neurons we recorded as having a higher firing rate during the ten seconds after speed transitions were responding to the unexpected change in the environment as a ’failure’ to match the speed of the wheel. We did find that a certain proportion of neurons were active even a minute past the transitions, but our experimental set-up—which was continuous running, without reward or pauses when the task was completed successfully—did not allow us to analyze whether the circuit recorded speed transitions as mistakes of some sort.

One of the most striking clinical features of Rett Syndrome is the loss of acquired skills in the second year of life. Acquired motor and language skills can disappear in a matter of days to weeks. At a behavioral level, the children seem unable to retain what they have previously learned or to learn new skills. Whatever roles MeCP2 has in early development (*54*), it seems to have additional functions in neuronal or synaptic maintenance (*11*). The period between 10 and 15 weeks of age is one of heightened plasticity, when we see elaboration of dendritic arbors and elevated expression of synaptic proteins; these processes seem to be profoundly disrupted in the absence of MeCP2 (*55, 56*). Despite these defects, the null mouse motor circuits showed plasticity in L2/3 (**Fig. 2**) and across layers (**Fig. 3**) in a similar pattern as in WT animals. The new synaptic connections initiated by null L2/3 and L5a neurons, however, tended not to last as long, which further supports the notion that MeCP2 is involved in neuronal maintenance. The number of these short-lived functional connections in null mice was over twice the number observed in wild-type animals (**Fig. 4B**), though this could be because the greater density of neurons on the null mice allowed us to record many more neurons. These observations call to mind an earlier study of the visual circuit in *Mecp2*-null mice. Noutel et al. found that the initial development of synapses between retinal ganglion cells and relay neurons in the dorsal lateral geniculate nucleus proceeded normally, but that synapses failed to be strengthened during the subsequent period of visual experience-dependent plasticity the *Mecp2*-null mice (*57*). A previous *ex vivo* study using glutamate uncaging and laser scanning photostimulation (LSPS) in M1 slices showed that knocking down MeCP2 reduces connectivity between M1 L2/3 and L3/5a neurons (*35*). Dani and Nelson (*36*) used paired recordings to reveal that there were fewer connections to L5 pyramidal neurons from the somatosensory cortex of *Mecp2*-null mice; moreover, individual connections were weaker. These results using slice recording align well with our findings that the connectivity of M1 pyramidal neurons is reduced in *Mecp2*-null mice.

The current study shows that two weeks of training reduced anxiety and extended the lifespan of symptomatic *Mecp2*-null mice by 20%. We had previously shown that deep brain stimulation (DBS) rescues hippocampal-dependent learning and memory in RTT mice (*18*) because it corrects hippocampal circuit abnormalities (*19*). DBS provides a very strong stimulus, however, around 130 Hz, which is considerably higher than the normal firing rate for neurons. Wheel-running is a much more ’physiological’ stimulus, and still is sufficient to alter circuit function and behavior of symptomatic *Mecp2*-null mice. While this manuscript was in revision, Achilly et al. published a study showing that exercise benefits female *Mecp2-*heterozygous mice who are not yet symptomatic (*58*). They also found some benefit for mice who were trained after becoming symptomatic, though they de-emphasize this result. Their mice did not appear to benefit in other realms, however, so they conclude that benefits of exercise are task-specific. We suspect that the broader benefits we observed came from our different training strategy. First, we used a forced exercise paradigm; otherwise, symptomatic *Mecp2* null male or heterozygous female mice are too sick to train. The fact that the head fixation prevented the mice from having to support their full body weight probably allowed them to train more intensively than they could on the accelerating rotarod. (Even our trained mice stopped running on the rotarod when they got tired; we suspect exercise fatigue (*23*) or apraxia would have been a limiting factor in the Achilly study, which used the rotarod for training.) Second, our circuit-level analyses suggest that mutant mice would learn even better with daily training than with the biweekly rotarod training regimen used in (*58*).

We also suspect that the additional benefits we observed (diminished anxiety, increased lifespan) involve motor learning as distinct from exercise *per se*. Exercise alters multiple neural circuits through stimulating BDNF and many other changes, as has been documented in many studies (*40, 44, 45, 59*), and the fact that rotarod training in (*58*) delayed the onset of disease symptoms in RTT mice speaks powerfully in favor of exercise. In favor of our hypothesis, however, we found a positive though weak correlation between extension of lifespan and improvement in forepaw coordination. It is worth noting in this context that Achilly et al. found that training in the Morris Water maze, which arguably involves more learning than exercise, led to improvements in the morphology and function of hippocampal neurons (*58*). Indeed, their Morris water maze training regimen (4 days per week) may have improved cognition because it works within the two-day window of functional connections we observed—if we assume that other *Mecp2*-deficient pyramidal cells behave in the same way as those in M1, which may not be the case. Regardless, the improved dendritic arborization they observed in hippocampal neurons from water-maze-trained mice is consistent with a hypothesis that learning may contribute benefits that are distinct from those of exercise.

There have been a few case reports and small studies showing Rett girls benefit from exercise (*60–64*), and the current study certainly confirms that these reports should not be dismissed as anecdotal. Early environmental enrichment in Rett syndrome mouse models have also reduced various behavior deficits (*44, 45, 65, 66*). This suggests that stimulation in general —in other words, exercise and learning of all sorts—is beneficial in Rett, as it is in healthy humans.

## Materials and Methods

### Animals

Mice were housed in an AAALAS-certified Level 3 facility on a 14-hour light cycle. Male *Camk2-cre; Mecp2^-/y^* (null) mice and female *Camk2-cre; Mecp2^-/+^* mice were obtained by breeding heterozygous female *Mecp2^+/-^* mice on the 129S6SvEvTac strain (*20*) (gift of Huda Zoghbi) to male *Camk2-cre* mice carrying the *Camk2-cre* allele on the C57/B6 strain obtained from Jackson Labs (JAX#005359).

All procedures to maintain and use these mice were approved by the Institutional Animal Care and Use Committee for George Washington University.

### Surgeries

At the age of 4 weeks, male mice were deeply anesthetized, then prepared for stereotaxic surgery. A 3-mm-wide hole was drilled through the skull over the motor cortex. The center of the craniotomy was ∼1.6 mm lateral to bregma (*67*). AAV/DJ-flex-GCaMP6m virus was injected into the forelimb area of the right motor cortex (1.5 mm lateral and 1.5 mm anterior to bregma, according to previous studies (*4*)) at two different depths (200 and 450 μm), to reach L2/3 and L5a neurons, respectively, via a 1.0-mm O.D. glass microneedle with a 10-20 micrometer diameter tip attached to a Nanoject microinjector pump. A glass coverslip was then placed over the exposed brain and sealed into the skull. The surgical site was closed using veterinary surgical glue. A 2-gram titanium headpost washer was attached to the head with “cold cure” denture material for later head fixation under a two-photon microscope.

To verify imaging depth and location of viral expression, three animals were perfused with 10% formaldehyde and the brains were sectioned. The depth for L5a was confirmed with three Rbp4-Cre mice as shown in **fig. S2a**.

### Chronic in vivo two-photon imaging

The experimental animal was lightly sedated with isoflurane and placed, head-fixed to the microscope, on a wheel, then allowed to wake up. The mouse was acclimated to the wheel in the free-running mode for 5 minutes, head-fixed under a Thorlabs Bergamo II two-photon microscope (Thorlabs, NJ) with high-speed piezo-Z optical-axis objective positioner and a Nikon 25X (NA 1.1) water-immersion objective lens (Nikon Instruments Inc., NY). Time-lapse images were acquired with Thorlabs software at 6-7 frames/sec/layer for 4630 frames/layer for 12 min (2 min for each speed for both speeding up and slowing down modes). The interval between the two modes at each trial was 2 min. The laser power was adjusted to the minimum necessary to achieve ideal fluorescence intensities during each imaging session. On average, about 20-30 mW of power arrives at the sample at layer (L) 2/3 and 120-160 mW of power at L5a. In WT mice (n=13), we imaged a total of 28,125 L2/3 neurons and 10,395 L5a neurons; in *Mecp2*-null mice (n=14) we imaged a total of 44,585 L2/3 neurons and 20,545 L5a neurons. The reason for the greater number of recorded neurons in the null mice is that they have smaller, more densely packed cells. The ratio of null to WT neurons recorded in this study is ∼1.47, which is very close to the 1.5 derived from an earlier study (*14*) which found that the cell density of L2/3 neurons in the somatosensory cortex of *Mecp2*-null mice after 8 weeks of age is about 120 neurons / 10^5^ μm^2^, while it is only about 80 neurons / 10^5^ μm^2^ in WT mice.

### Set-up for studying motor skill learning over a two-week period

Beginning four weeks after surgery, experimental animals were trained daily for 14 consecutive days on a computerized wheel system customized by Delta Commercial Vision (NJ, USA) while two-photon imaging was performed simultaneously (**Fig. 2A**). The six-inch-diameter foam wheel is connected to a motor controlled by an Adreno chip connected to a computer via a USB cable, so that the experimenter can control the speed of the wheel rotation. There are five speeds: 0, 15, 30, 45, 60 mm/sec, and the experiment maintained each speed for 2 min, either in ascending or descending order.

Before the start of each trial, the experimental animal was put on the treadmill with the head fixed at a comfortable height under the two-photon microscope. ThorCam software controlled the videotaping of the mouse at 71.5 Hz via a high-speed camera (Thorlabs, NJ) and simultaneously triggered the ThorImage software (Thorlabs, NJ) to start two-photon imaging and the digital command that controls the treadmill. The experimental task is to learn to smoothly and quickly adapt to running at different speeds.

Each trial comprised three sessions. “Session 1” was 12 min in total. It started at speed 0 (rest) and increased by 15 mm/sec every 2 min up to 60 mm/sec, followed by a 2-min rest period. “Session 2” was a 2-min interval for the animal to rest and for the experimenter to save the data. “Session 3,” also 12 min in total, started at speed 0, jumped to 60 mm/sec and remained there for 2 min, then decreased by 15 mm/sec every 2 min down to 0 mm/sec. There was one trial per day for two weeks.

### Motion analysis

We detected paw positions with an efficient pose estimation method called DeepLabCut which is based on transfer learning with deep neural networks. We chose 3 days from the 14 days of experimental data and chose the period at speed 60 for training. We next used 5 body markers to track forelimb movement. We tracked 5 points: the center of the two forepaws, two front knees, and the center of the chest to form a “skeleton.” For every training image, we dragged each skeleton point to the appropriate body part, and the program saved the label positions automatically. With limited training data, we achieved a satisfying model to run on all movies and detect paw locations. Then we saved the coordinates of all the body markers and used the Butterworth filter to process the raw data with order 2 to reduce the influence of error.

To get the paw location value (d), we first chose the median of the paw coordinates as the zero point. We gave forward paw positions positive position values and backward paw positions negative position values.

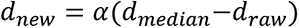

Where *d*_*raw*_ is the original distance between the forepaw’s location and the lower right corner, *d*_*median*_ is the median of the distance and *α* is a parameter used to convert pixels into millimeter. According to the location of the paw, the start and endpoint will be detected by an algorithm.

As shown in **Fig. 1C**, the x- and y-axis represent the direction of paw movements along the horizontal and vertical planes, respectively. Stride was represented by the rising phase of a spike in X; a complete spike in the paw coordination index was defined as the negative value of the Pearson correlation coefficient of the two paw location traces.

### Calcium image processing

Calcium images were processed and analyzed using custom scripts in Python based on CalmAn (*68, 69*), a toolbox for large-scale calcium imaging data analysis and behavioral analysis. This toolbox provides fast and scalable algorithms for motion correction, source extraction, spike deconvolution, and component registration across multiple days.

Source extraction was performed using the online algorithms (*69*) by adapting constrained nonnegative matrix factorization approaches (*68, 70*). For the prediction of spikes from fluorescence traces, we applied unsupervised learning sparse non-negative deconvolution (*80, 83*) and near-online OASIS algorithm (*72*) to our raw calcium signals. To align calcium recordings collected from speed-increasing and -decreasing modes, we first artificially stacked all calcium videos collected in one trial for the same mouse and then CaImAn performed motion correction using the NoRMC algorithm (*73*), which corrects non-rigid motion artifacts by estimating motion vectors with subpixel resolution over a set of overlapping patches within the field of view (FOV). The width of event rate distribution is calculated as full width at half maximum of the distribution by the function scipy.signal.peak_widths in the scipy package in Python.

### Detecting the same neuron across trials

Although we imaged the same area of the motor cortex for each experiment, little shifts do occur in the field of view (FOV) between trials. Thus, both the total number and precise location of detected neurons can vary somewhat across trials. To accommodate this slight inconsistency and enable direct comparison between trials for each mouse, we aligned the images between days and picked the same neurons throughout 14 trials. The basic workflow is shown in **fig. S6A**. First, we constructed an image of the distribution of neurons based on their pixel location.

Instead of representing a neuron in a single pixel, we used a bivariate normal distribution peak centered at its location with maximum height 1.0 and STD as 5, which is around the radius of real neurons in the frame. The reconstructed image has the same size as the raw frame (512 x 512), as shown in **fig. S6A**. For each sample (from each trial), we used the detected neuron from Trial 1 as a reference and aligned all 14 images (including subsequent trials). To do this, we expanded the image of Trial 1 slightly so that neurons that would otherwise appear slightly ‘displaced’ from the first image became aligned with the selected neuron; we created a border around the image with a width of approximately 80 pixels to accommodate the maximum shift we observed between Day 1 and any other trial (and prevent the program from having nothing to compare at the edges of the FOV). For each map, we calculated the correlation score of the two-image matrix (Day 1 x Day *N*), resulting in a correlation matrix of 81 x 81 (**fig. S6A**, step 5).

Correlation matrices have a size of 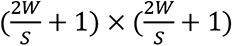; the shift (Δ*x,* Δ*y*) with the highest correlation score indicates the degree of shift of the image from Day 1 to the present image. A desired correlation matrix should be sparse, i.e., except for a very small region with high correlation score, most parts should be nearly zero.

Given a known degree of shift between two trials, it becomes straightforward to map the same neuron by correcting the location of neurons from the subsequent trial and matching to Day 1. Considering the various shapes and sizes of neurons, we allowed for a difference of 5 pixels when matching. Thus, for each neuron *N*_*i*_(*x*_*i*_,*y*_*i*_) at Day 1, we searched for the same neuron on the subsequent trial image, and if there was a neuron within 5 pixels of (*x*_*i*_,*y*_*i*_), we assumed it was the same neuron; otherwise, we recorded no corresponding neuron on that trial’s image. If there was more than one neuron within 5 pixels, we chose the one closest to the original *N*_*i*_. **fig S6A**, step 6 shows examples of matched neurons over two different days. On average, 20% of the total active neurons detected were consistently recorded each day throughout 12days.

### Identification of neurons responsive to speed transitions

We identified neurons particularly responsive to speed transitions based on their response each time the computerized wheel transitioned from one speed to another, in either session, following the steps below:

(1) For any neuron *k*, we calculated the similarity between the original calcium trace (Δ*F*/*F*) *c*_*k*_ and the speed transition vector *s*, where *s*_*t*_ = 1 if time *t* was within 15 seconds of every speed transition, otherwise *s*_*t*_ = 0. The similarity was defined as the normalized inner product of two vectors, 2*s* ⋅ *c*_*k*_/(|*s*|^2^ + |*c*_*k*_|^2^). The value of the similarity was between 0 and 1. A value of 1 meant these two vectors were identically distributed while a value of 0 meant these two vectors were completely different.
(2) We then randomly shuffled the speed transition vector *s* 5000 times and calculated the similarity between *c*_*k*_ and *s*shuffled to generate a similarity distribution representing the similarity histogram predicted by chance for neuron *k*.
(3) Neuron *k* was defined as a “transition-active neuron” if its actual similarity was greater than 99.95% of the distribution, representing a significantly higher possibility of a positive correlation between neuronal activity and speed transition.

### Behavioral testing

After two weeks’ training on the variable-speed wheel, we tested mice on the open field, two-tiered ladder, accelerating rotarod, elevated plus maze, and grip strength tests with age-matched littermates who had not been trained (naïve mice). For the open field test, a mouse was released into the center of the square open field arena, then allowed to move freely for 10 minutes without interruption or interference. A fast-speed camera on top of the arena captured video of the mouse during the test. The videos were analyzed by software TopScan (Clever Sys., Inc., VA, USA).

A ladder-walking behavior analyzing system (Clever Sys., Inc., VA, USA) was used for the ladder test. This is a horizontal ladder composed of 2×37 rungs. The rungs are spaced 15 mm apart and alternate heights between high and low, so that the mice prefer to walk on the high rungs. Steps on low rungs are considered miss-steps. Thus, fewer miss-steps means better motor coordination. There is one shelter box at each end of the ladder. In the beginning of the test, one mouse is released into the left shelter box. Then, 5 seconds after the mouse leaves the box to walk the ladder, a light is turned on in the shelter box. A fan is turned on after another 5 seconds to further encourage the mouse to walk the ladder rather than return to the safety of the shelter. Walking from one end of the ladder to the other is defined as one trial. Each mouse performed twenty trials, during which we measured the time needed to cross the ladder and the number of correct steps (steps between adjacent high rungs) and irregular steps (steps from high rung to low rung, from low rung to low rung, crossing boundary to rungs on the other side, and backward steps).

The accelerating rotarod assay was performed as previously described (*74*). *Mecp2*-null mice and controls were placed on an accelerating rotarod apparatus (IITC Inc.) for 4 trials in one day with a 15-minute rest interval between trials. Each trial lasted for a maximum of 5 min, during which the rod accelerated linearly from 4 to 40 rpm. For each trial we recorded the latency to fall (i.e., the amount of time it took for each mouse to fall from the rod) and noted when the mice first stopped locomoting by holding the rotating rod.

The Elevated Plus Maze assay was performed as previously described (*75*). The elevated plus maze apparatus consists of a plus-shaped platform that is elevated above the floor. Two opposite arms of the maze are walled, whereas the other two arms are open with a small threshold along the sides. Mice were placed in the center part of the maze facing one of the two open arms. The number of entries and the amount of time the animal spent in the open and closed arms were scored with Topscan behavioral data acquisition software (CleverSys, Reston, VA, USA).

For the grip strength test, each mouse was tested twice on two adjacent days by following the procedure as previously described (*76*). Each mouse was placed on a grid and exercised by pulling its tail without the mouse losing its grip. The action was repeated for 20 paw grips. The following five grips were measured by a dynamometer (Chatillon Digital Force Gauge, DFIS 2, Columbus Instruments, Columbus, OH). The maximal force applied to the dynamometer while on the grid was recorded. The maximum value of the two tests was used in **fig. S12A**. We also compared the mean value of two days’ tests and got same results.

## Statistics

Linear mixed models were used to obtain p-values. All response types are assumed to be Gaussian. Fixed effects in the linear regression model are given by (i) a factor variable that represents the trial number (i.e., day of the experiment), and (ii) the mouse type (i.e., wild-type or *Mecp2*-null). The presented p-values are for the latter covariate. In addition, separate random effects for each mouse were fit, and an AR(1) correlation model was used to model the temporal correlation across the trials. We used R (R Core Team, 2019, https://www.R-project.org) and the R package “lmerTest” (*77*) or SPSS Version 26 (IBM Inc., Chicago, IL, USA) for fitting the models and performing inferences. In Fig. 3A and B, p-values are obtained under Wilcoxon signed-rank test by SPSS.

## Acknowledgments

We would like to thank Dr. Huda Zoghbi at Baylor College of Medicine for providing the *Mecp2* null mice. We also thank Ethan Jin for significant input in data analysis and V.L. Brandt for many probing discussions and her commitment to clarity in editing the manuscript.

## Author contributions

Y.Y., P.X., L.K.and H.L. designed and performed the experiments. S.S. came up with the idea of the computerized wheel and supervised the initial experiments. L.K. performed the grip strength test. Y.Y., Z.L., J.S., H.D. and L.C. analyzed data. Y.Y., P.X., Z.L., J.S., H. D., L.C., R.T.A., L.K., C.Z., and H.L. reviewed and interpreted data. Y.Y., Z.L., H.D., L. K., Z.C., J.S., and H.L. wrote the manuscript with critical input from P.X., E.B., R.T.A., C.Z and S.S.

## Data availability statement

All data needed to evaluate the conclusions in the paper are present in the paper and/or the Supplementary Materials.

---

**Video 1.** A WT mouse running on the wheel at speed 60 mm/sec on Days 2 and 10.

**Video 2.** A *Mecp2*-null mouse running on the wheel at speed 60 mm/sec on Days 2 and 10.

**Video 3**. A naïve *Mecp2*-null mouse running on the ladder.

**Video 4**. A *Mecp2*-null mouse running on the ladder two days after the end of the two-week motor training.

## Supplemental information

**Fig. S1.**
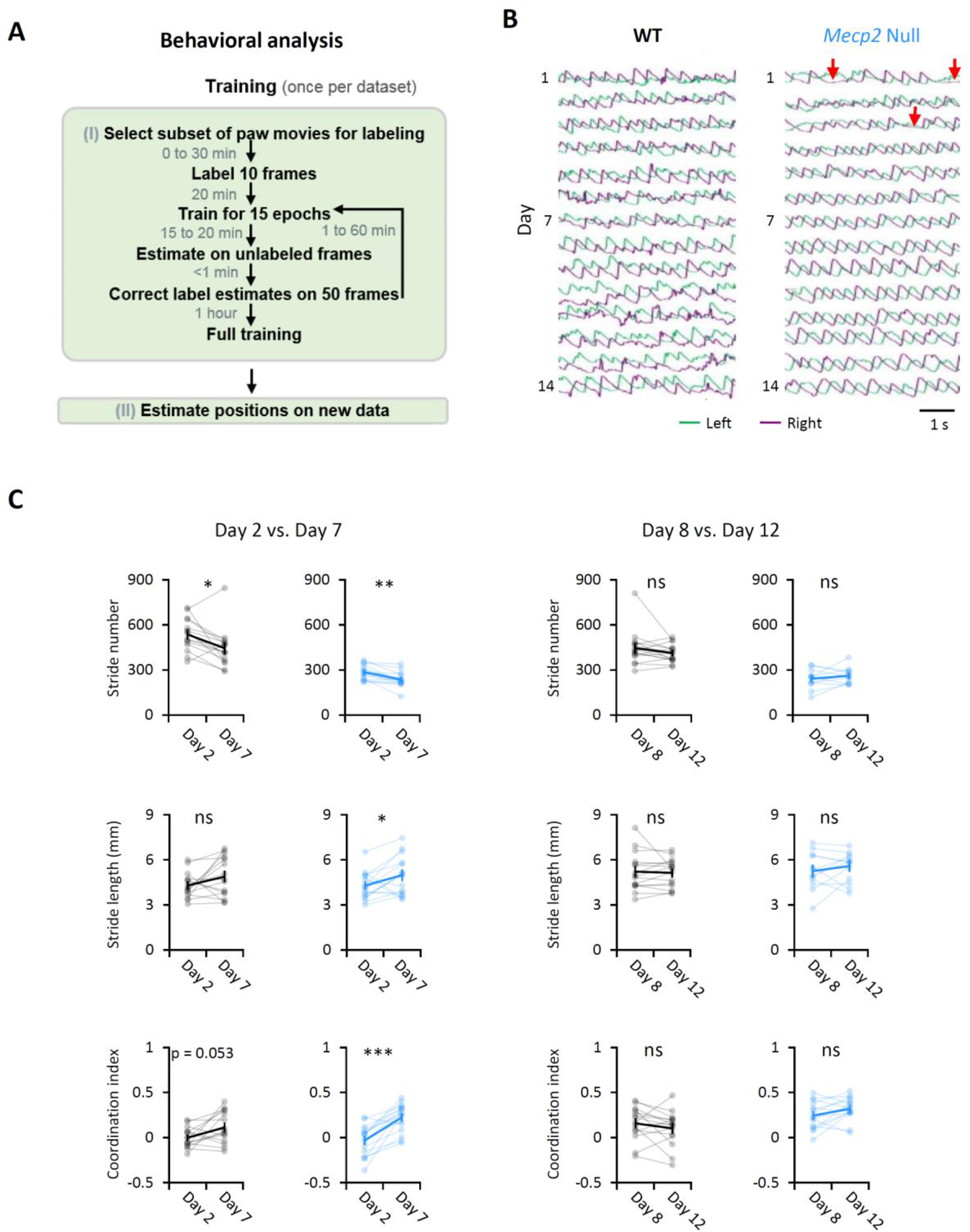
Analysis of forepaw movement during running and degree of motor improvement during the first week vs. second week of training. (**A**) The workflow of a machine learning approach, DeepLabCut method, to track forepaw location. (**B**) Raw traces of forepaw locations across 14 days at speed 60 mm/s from one sample mouse in each group. (**C**) Comparison between changes in stride and coordination during the first and second week of training show greater improvement during the learning phase than the consolidation phase. WT: n=12 mice; *Mecp2*-null, n=14 mice. ns, not significant, * *P* < 0.05, ** *P* < 0.01, Wilcoxon signed-rank test.

**Fig. S2.**
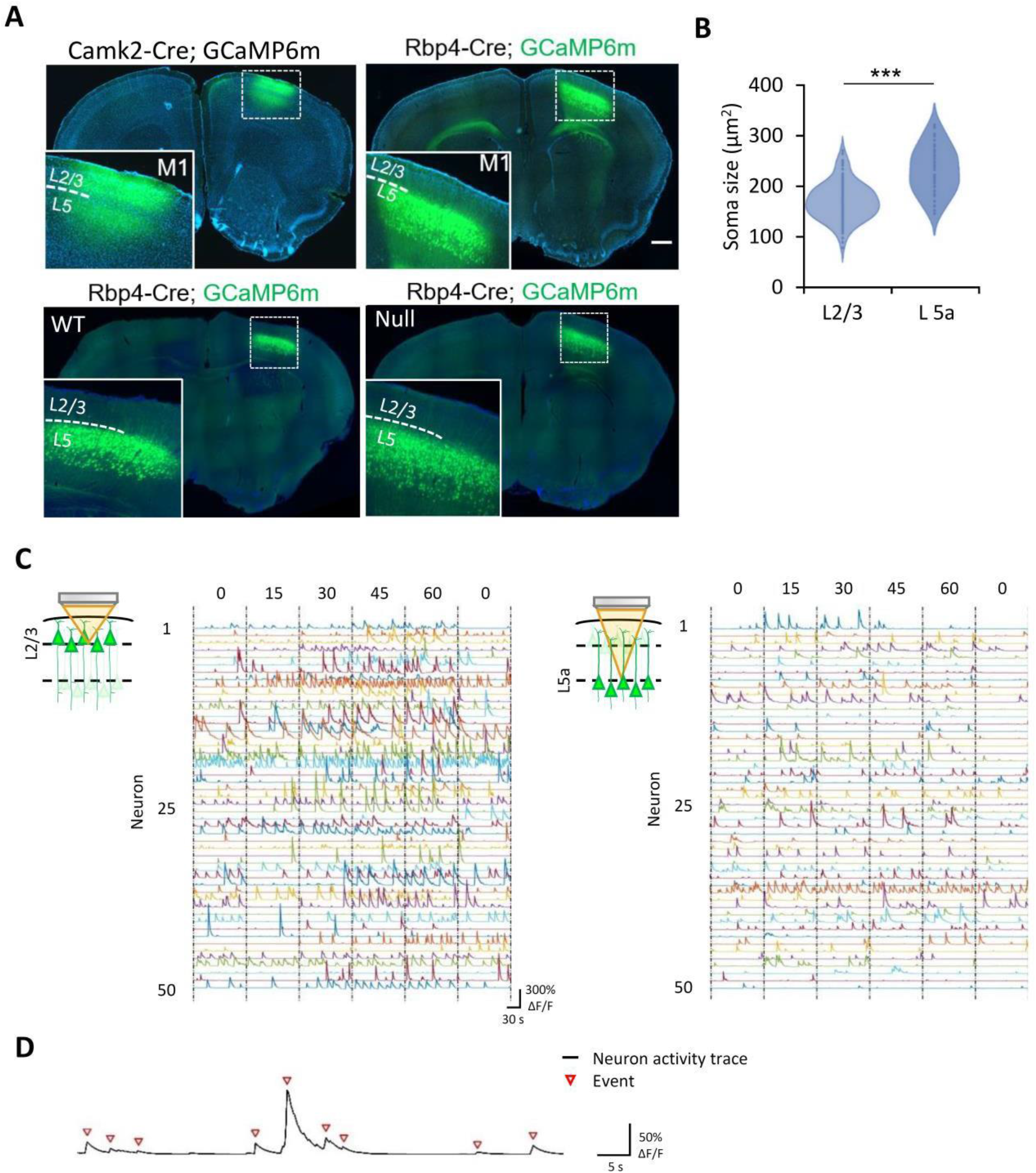
Confirmation of imaging location and processing of imaging data. (**A**) Images of coronal sections showing expression of GCaMP6m in the area M1. Magnification of the boxed area is shown in the lower left corner of the larger images. Top, Rbp4-Cre-induced expression confirms the depth of L5a isaround 400-450 µm. Bottom, no noticeable difference of the depth of L5a between the WT and *Mecp2*- null mouse. Scale: 500 µm. (**B**) Violin plot with the addition of a rotated kernel density plot on each side comparing the soma size of detected neurons in L2/3 and L5a. *** *P* < 0.001 (Rank sum test) (**C**) Pre- processed sample traces of calcium dynamics in 50 neurons in L2/3 (left) and L5a (right) of one mouse during increase-speed mode. (**D**) Denoised neuronal activity trace; red triangles point to events.

**Fig. S3.**
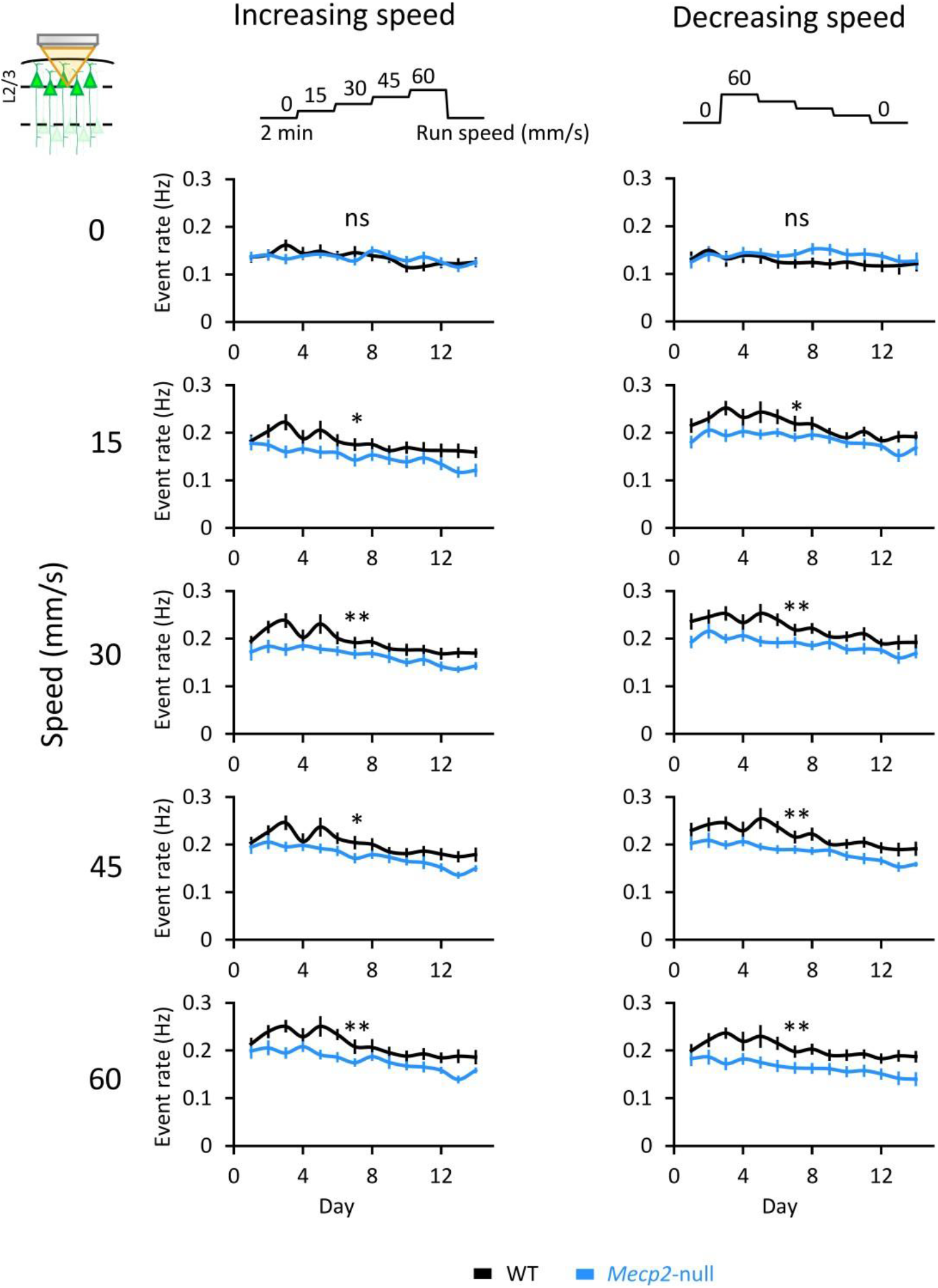
Comparison of firing rates at different speeds in M1 area L2/3. Summary of event rates of WT and null neurons in L2/3 in session 1 (left, speeding-up mode) and session 2 (right, slowing-down mode) at each speed. WT: n=13 mice; Null, n=14 mice. Error bars represent mean ± SE. ns, not significant; * *P* < 0.05; ** *P* < 0.01, RM-ANOVA test.

**Fig. S4.**
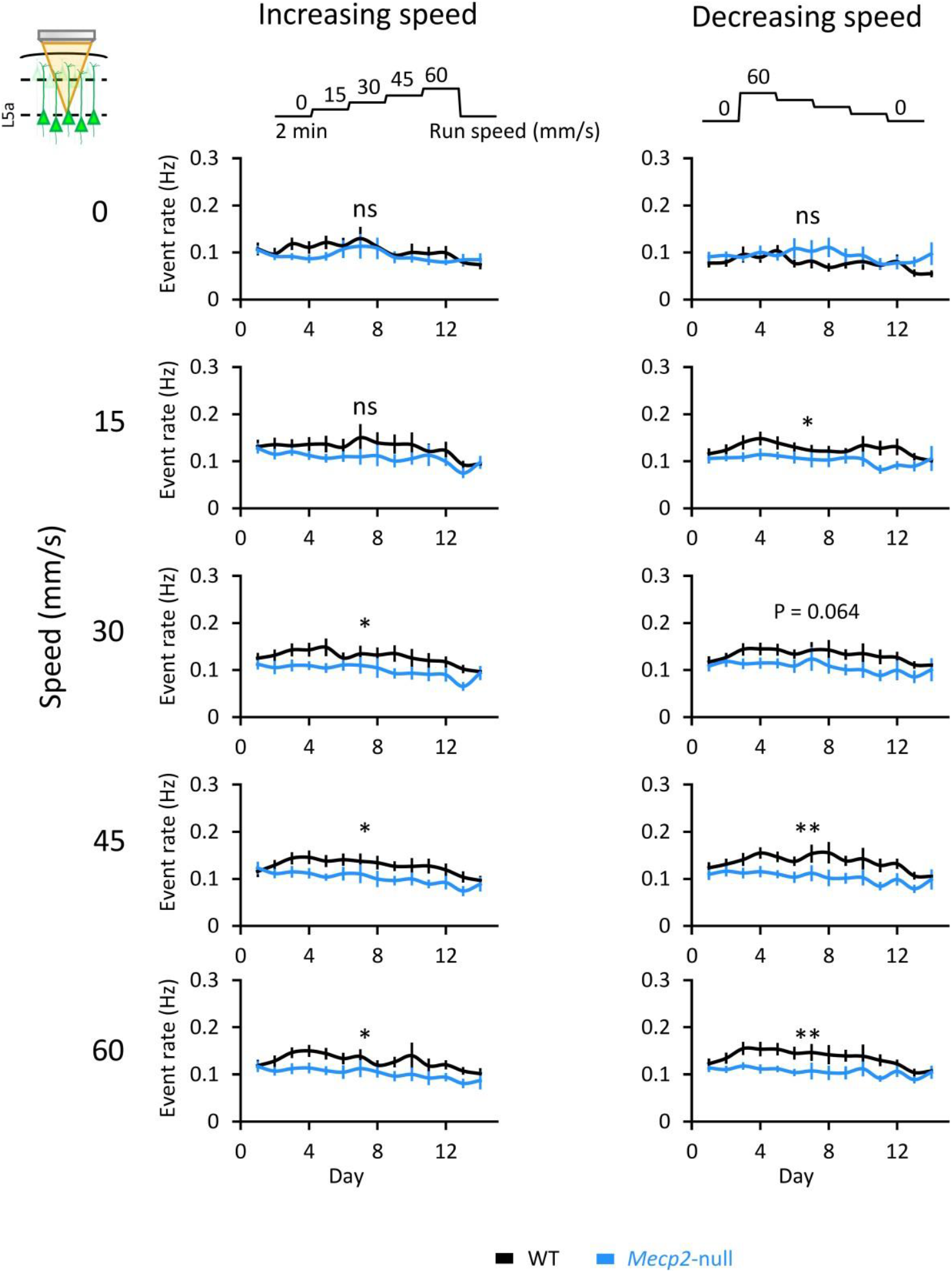
Comparison of L5a firing rates at different speeds. Summary of event rates of WT and Nullneurons in L5a in session 1 (left, speeding-up mode) and session 2 (right, slowing-down mode) at each speed. WT: n=11 mice; Null, n=13 mice. Error bars represent mean ± SE. * *P* < 0.05; ** *P* < 0.01, RM-ANOVA test.

**Fig. S5.**
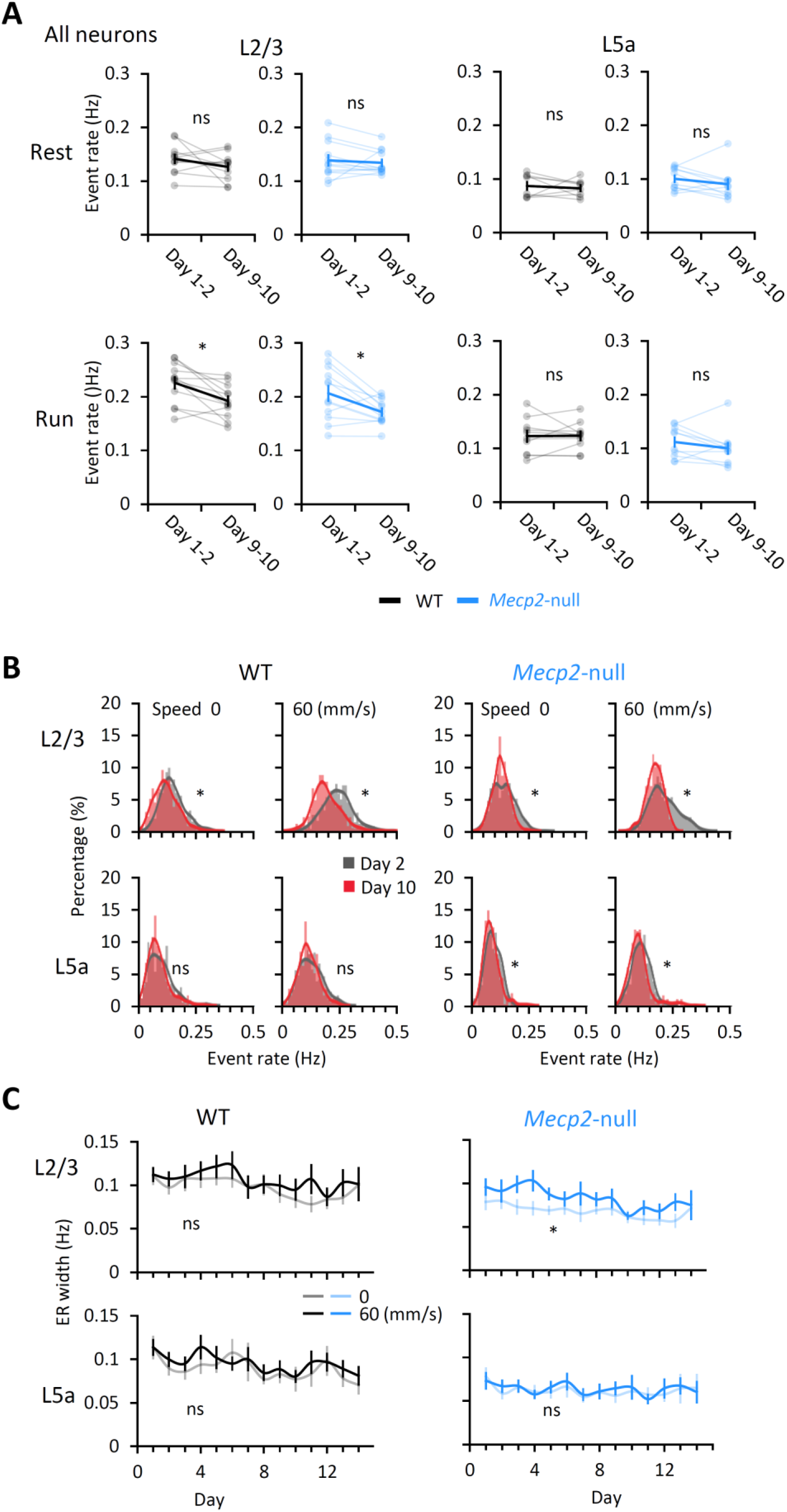
Evolution of event rates over the course of learning. (**A**) Mean event rates of neurons in L2/3 and L5a during rest and running at 60 mm/sec block. Light lines connect data from an individual mouse. Dark lines connect the averaged data from all mice. L2/3 WT: n=11 mice; L2/3 *Mecp2*-null, n=12 mice. L5a WT: n=8 mice; L5a *Mecp2*-null, n=10 mice. Error bars represent mean ± SE. ns, not significant, * *P* < 0.05, Wilcoxon signed-rank test. (**B**) Distribution of neuronal event rates of all mice on days 2 and 10. Motor learning narrows the distribution and lowers the rates over time. * *P* < 0.0001, Kolmogorov–Smirnov test. (**C**) The width of the event rate (ER) distribution in **B**, calculated as full width at half maximum of the amplitude of the distribution curve. L2/3 WT: n=13; L2/3 *Mecp2*- null: n=14; L5a WT: n=11; L5a *Mecp2*-null: n=13. * *P* < 0.05, ** *P* < 0.01, RM-ANOVA test.

**Fig. S6.**
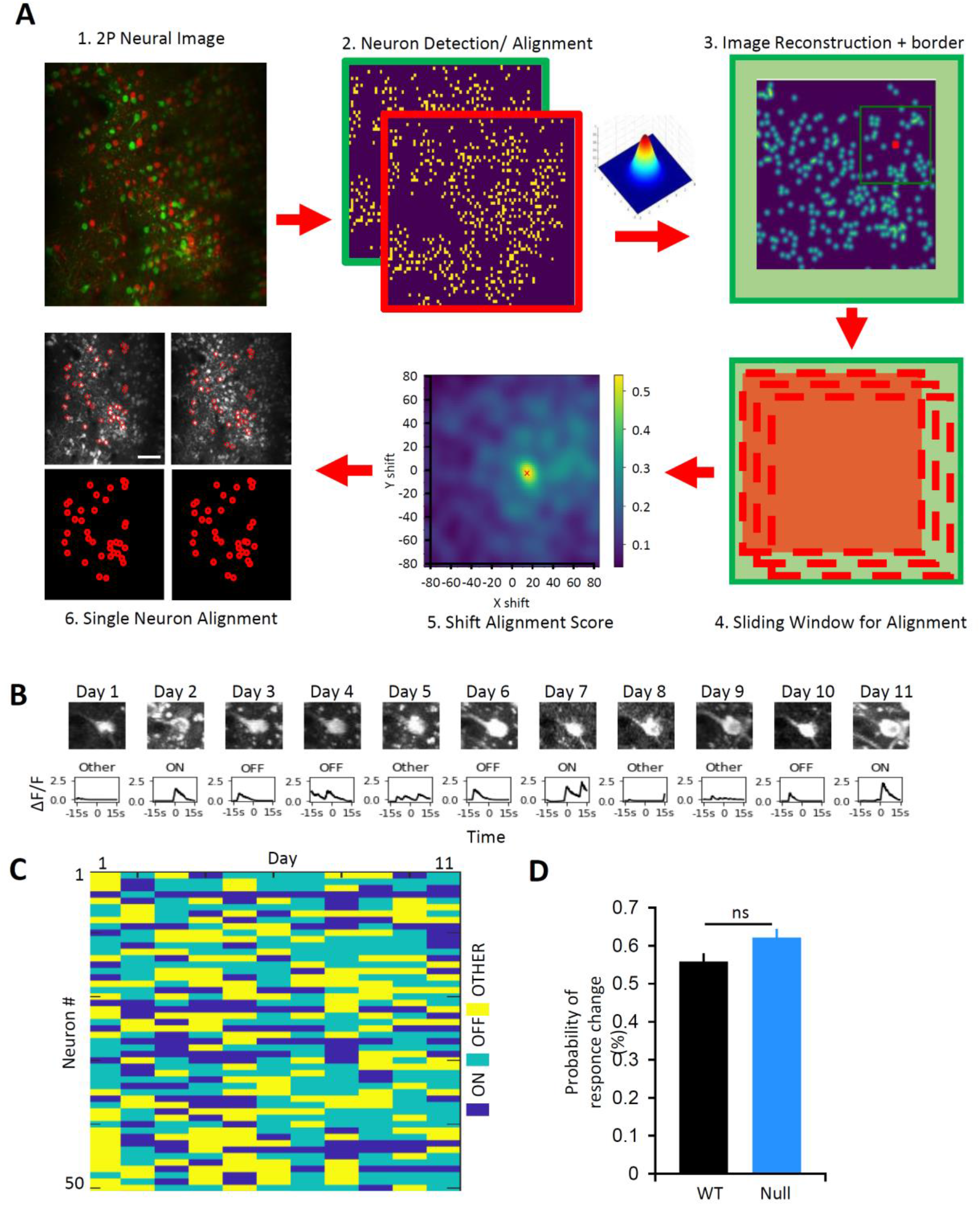
Behavior of one neuron over eleven days. (**A**) Schematic of Global Alignment method (see Methods). About 20% of neurons can be followed over 12 days. (**B**) A representative neuron that showed different responses to the transition from rest to 15 mm/s (bottom row) over 11 different days. (**C**) The responses of 50 individual neurons to a transition from rest to 15 mm/s on each day over 11days. (**D**) Probability of response type changes between two adjacent days in WT and *Mecp2*-null mice. WT, n=7 mice; *Mecp2*-null, n=5 mice. ns, not significant, Two-tailed t-test.

**Fig. S7.**
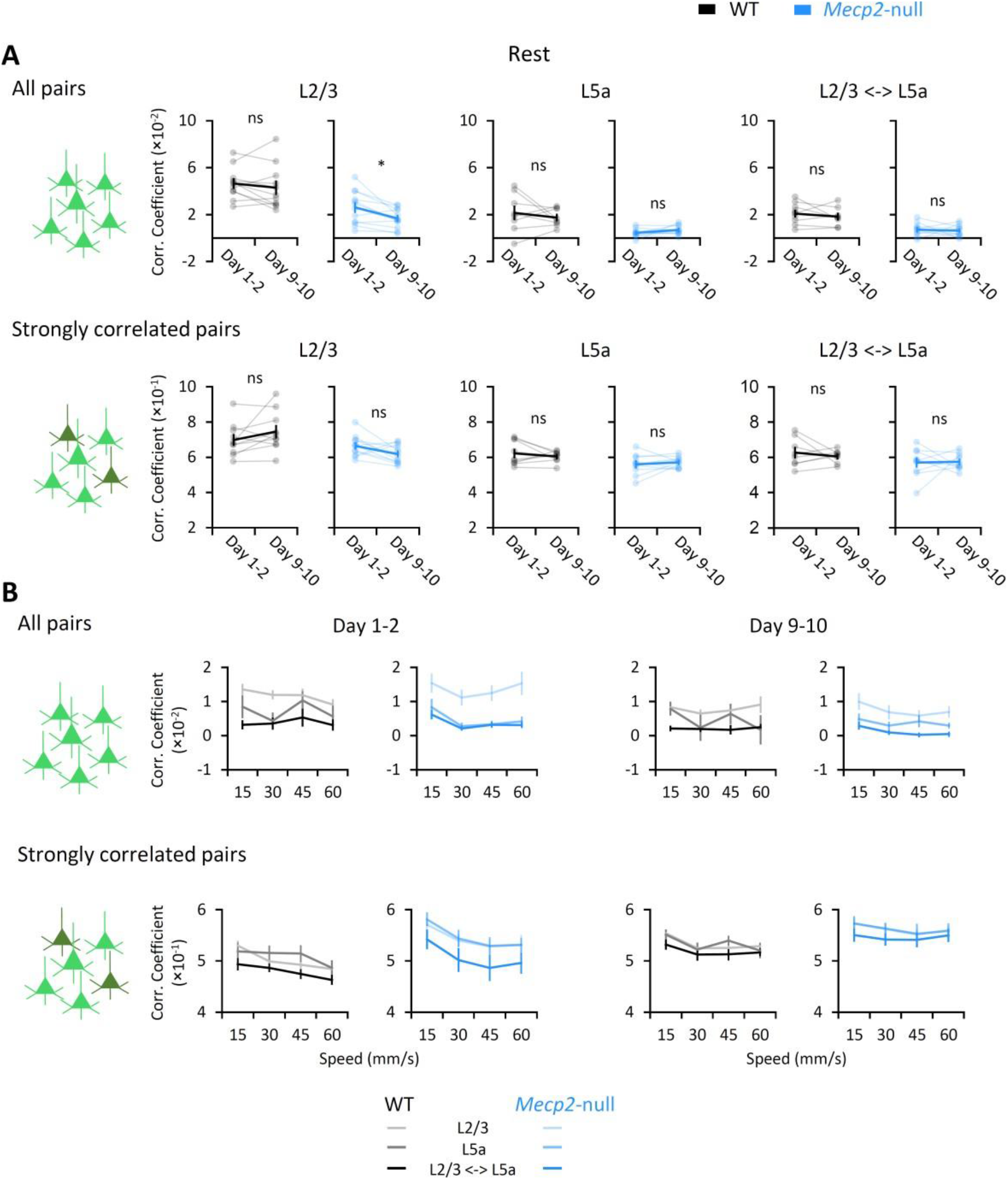
Evolution of functional connectivity at rest and running. (**A**) Mean Pearson correlation coefficient (PCC) between pairs of neurons within L2/3, within L5a, and across both layers during rest (top: all pairs of correlation; bottom: strongly correlated pairs, defined by a PCC > twice the standard deviation). Light lines connect data from an individual mouse; dark lines show the average from all mice. L2/3 WT, n=11 mice; L2/3 *Mecp2*-null, n=12 mice; L5a WT,n=8 mice; L5a *Mecp2*-null, n=10 mice. Across both layers, WT, n=8 mice; *Mecp2*-null, n=10 mice, Error bars represent mean ± SE. * *P* < 0.05, Wilcoxon signed-rank test. (**B**) Average PCC at each speed from all the mice of each group (top: all pairs; bottom: strongly correlated pairs).

**Fig. S8.**
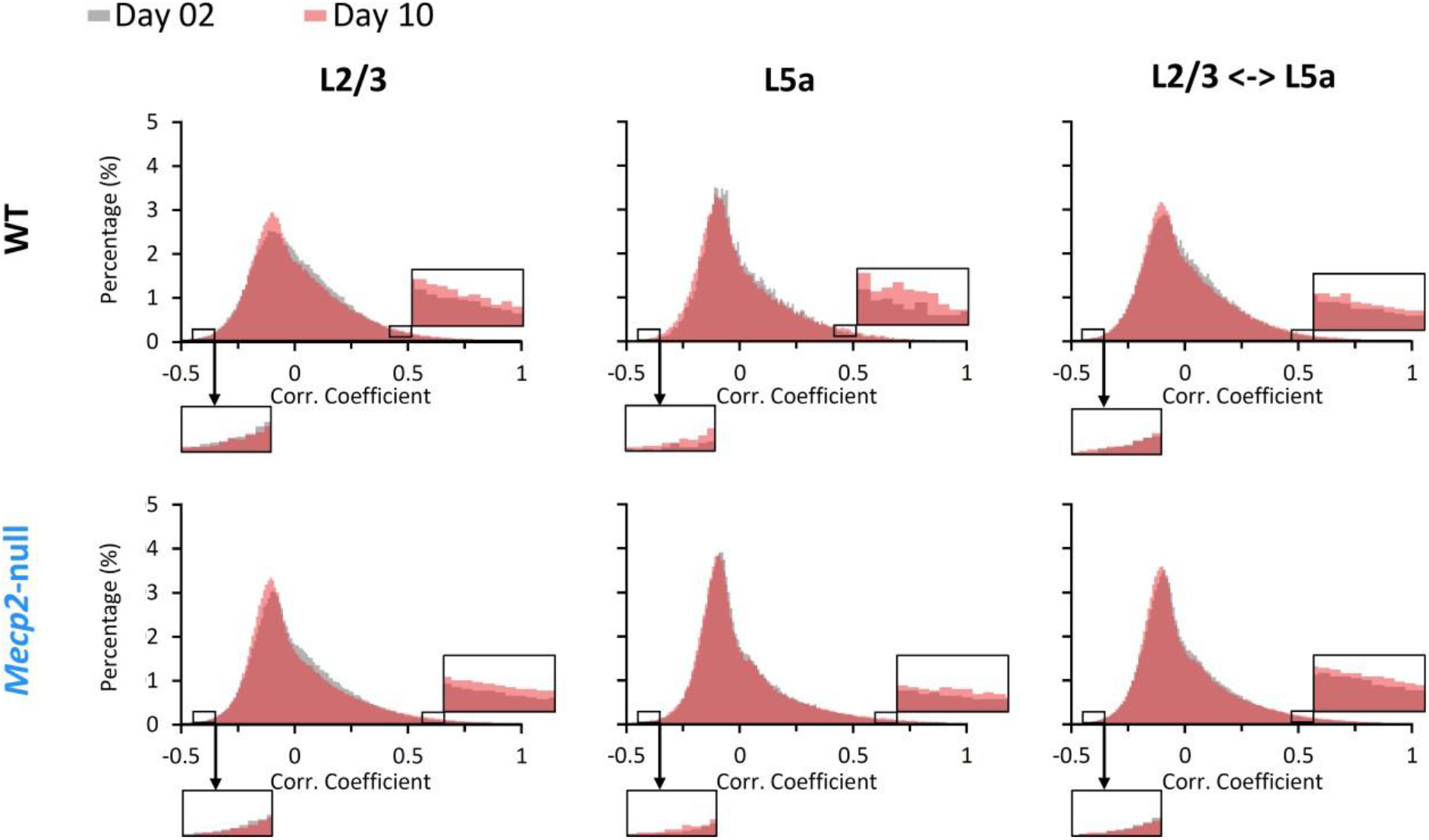
The distributions of Pearson’s correlation coefficients of functional neuronal pairs are very wide, reflecting the dynamism of M1. L2/3 WT, n=11 mice;L2/3 *Mecp2*-null, n=12 mice; L5a WT, n=8 mice; L5a *Mecp2*-null, n=10 mice (we excluded L5a data from mice from whom we detected fewer than 20 neurons in L5a). Across both layers, WT, n=8 mice; *Mecp2*-null, n=10 mice.

**Fig. S9.**
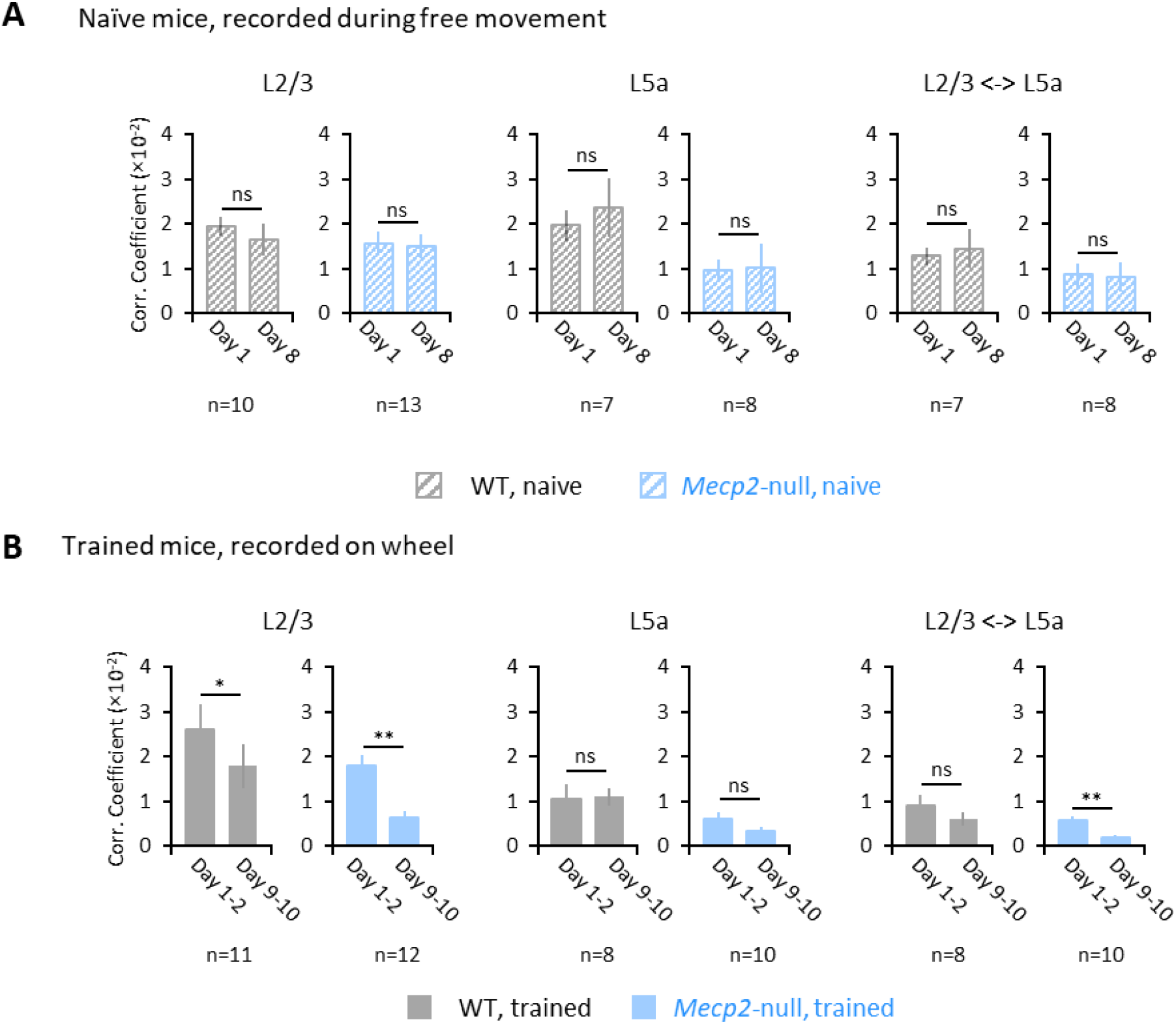
Circuit dynamics in naïve mice during free-running did not change over the course of a week, but the circuit in trained mice did. (**A**) Comparison of averaged correlation coefficients during free-run mode between days 1 and 8 in naive WT and *Mecp2*-null mice. There is no difference between days. (**B**) Comparison of averaged correlation coefficients during free-run mode between days 1-2 and 9-10 in trained WT and *Mecp2*-null mice. This indicates that the changes we observe in the M1 circuit over the course of learning do not just correlate with the behavioral improvements, but have a causal relationship to those improvements. Error bars represent mean ± SE. ns, not significant; * *P* < 0.05, ** *P* < 0.01, Wilcoxon signed-rank test.

**Fig. S10.**
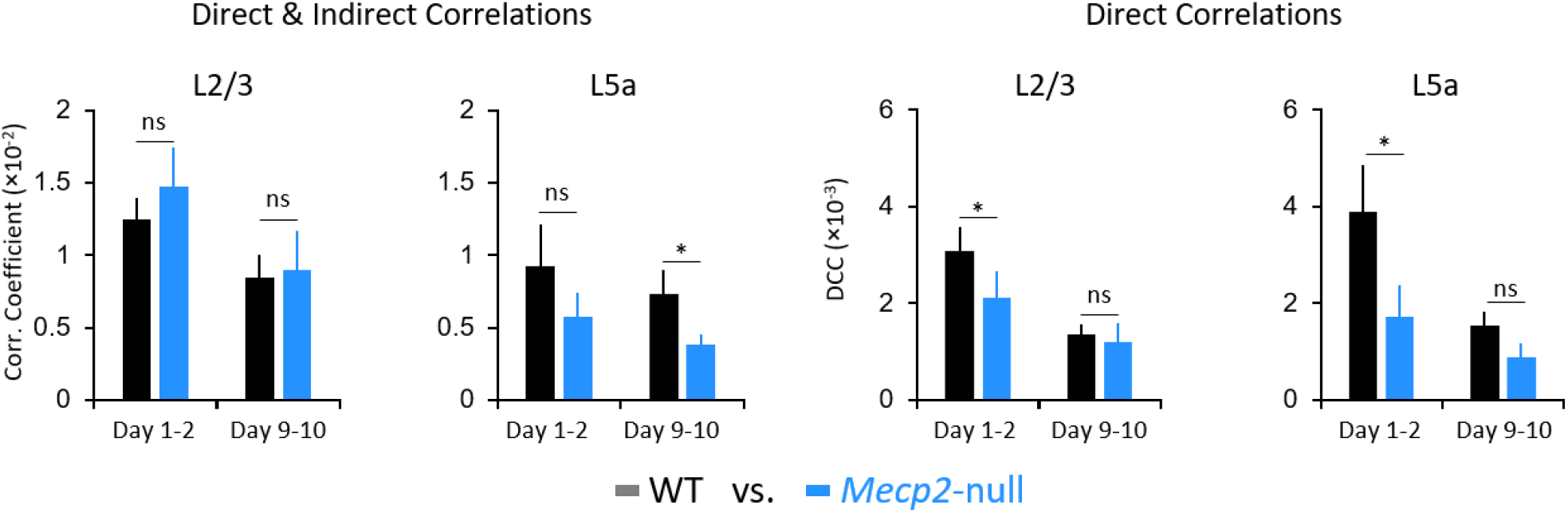
Cross-group comparison of Pearson’s correlation coefficient. Comparison of averaged correlation coefficients between WT and *Mecp2*-null mice during days 1-2 (learning) and days 9-10 (consolidation). The same data set was used as in Fig 3. L2/3 WT, n=11 mice; L2/3 Mecp2-null, n=12mice; L5a WT, n=8 mice; L5a Mecp2-null, n=10 mice. Error bars represent mean ± SE. ns, not significant; * *P* < 0.05, Mann-Whitney U Test.

**Fig. S11.**
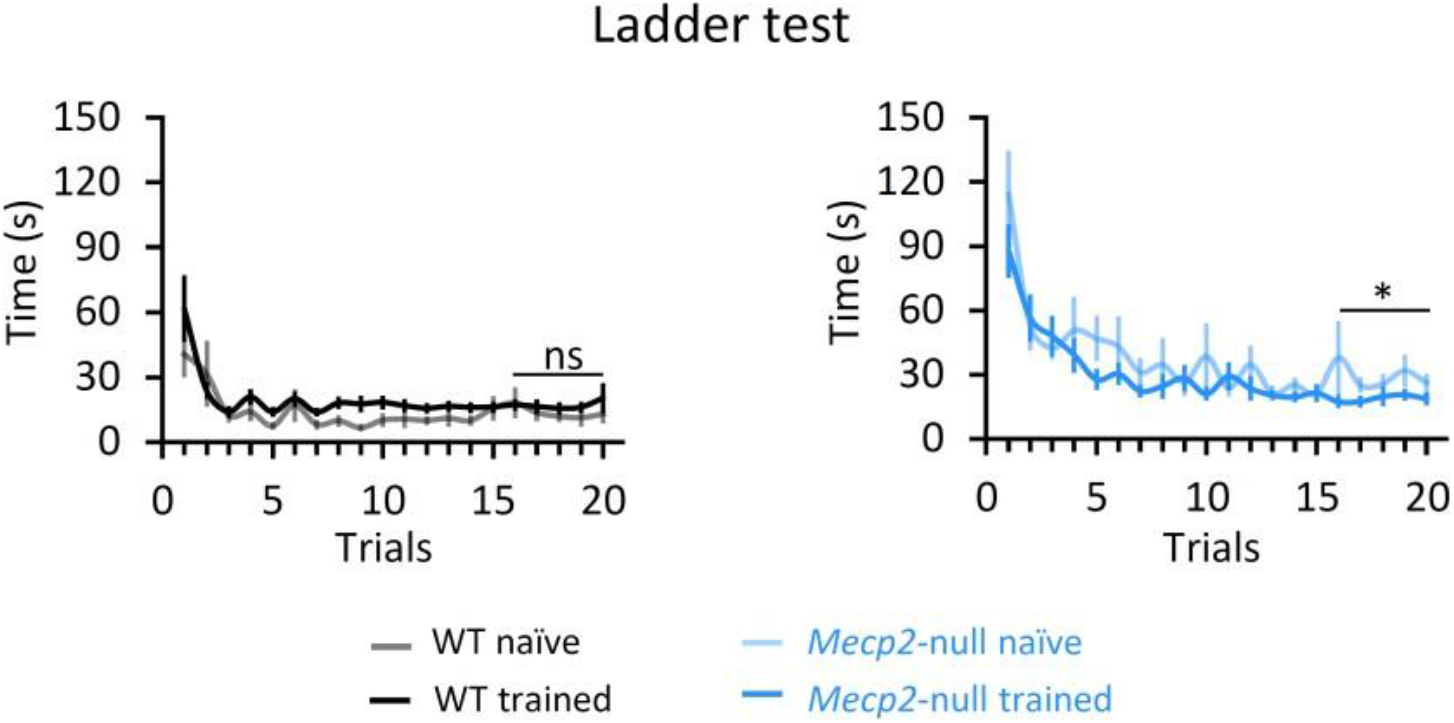
Wheel-speed-training improved the performance of *Mecp2* null mice on the ladder test. Graphs show the duration of time wild-type and null mice needed to cross the ladder from one end to the other over the course of 20 trials. The less time spent on the ladder, the better the mouse’s performance. Naive: WT: n=15 mice; *Mecp2*-null, n=17 mice; Trained: WT: n=19 mice; *Mecp2*-null, n=17 mice. Error bars represent mean ± SE. ns, not significant, * P< 0.05, RM-ANOVA test.

**Fig. S12.**
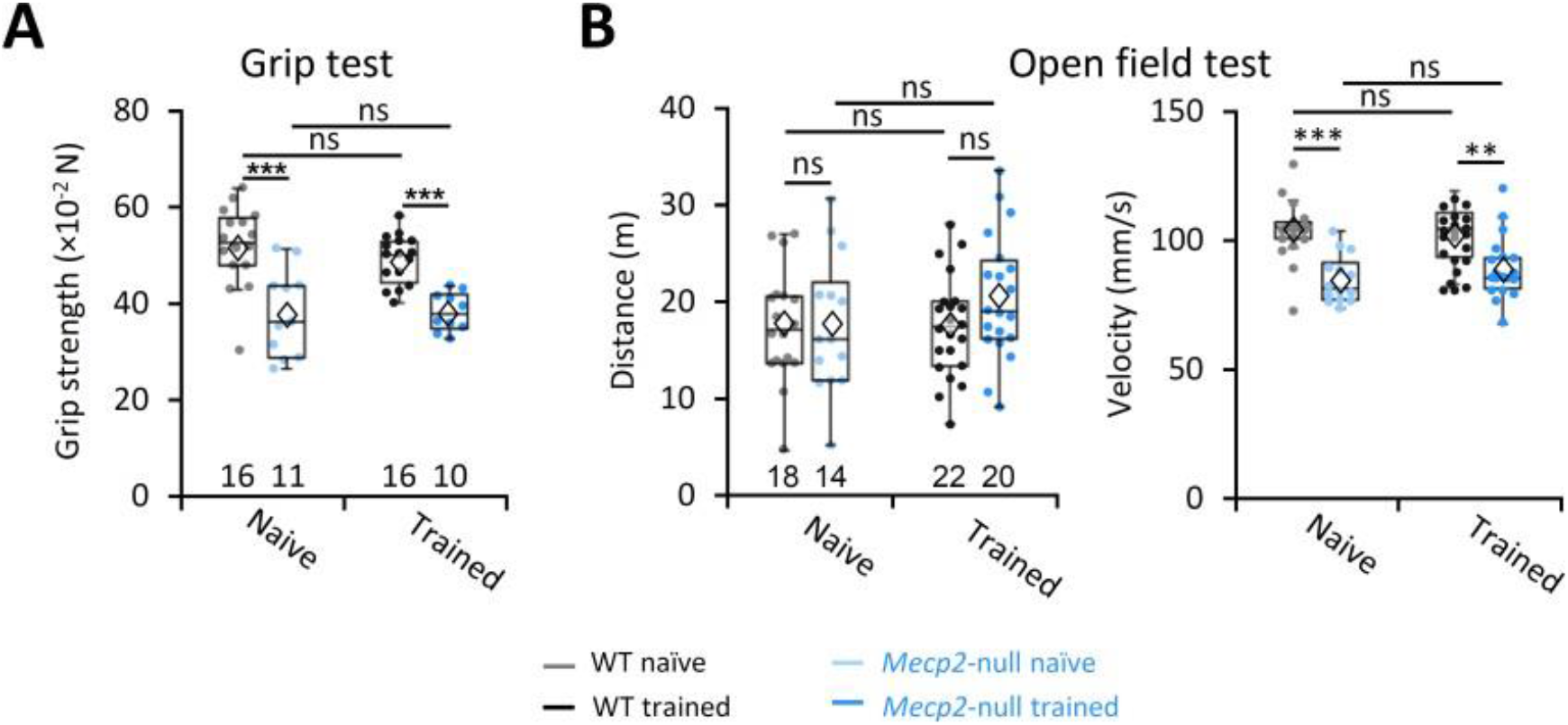
Motor training did not improve strength on the grip test or change parameters of physical performance on the open field test. (**A**) Summary of the grip strength of front limbs. Naive: WT: n=16 mice; *Mecp2*-null, n=11 mice; Trained: WT: n=16 mice; *Mecp2*-null, n=10 mice. Error bars represent mean ± SE. ns, not significant, * *P* < 0.05, ** *P* < 0.01, *** *P* < 0.001, Two-way ANOVA test with Sidak’s Post-Hoc. Diamond symbol represents the mean value. (**B**) Distance traveled in the open field testwas not affected by genotype; the null mice were slower than WT in moving around, regardless of training. The number of animals in each group is indicated beneath each bar. Naive: WT: n=18 mice; *Mecp2*-null, n=14 mice; Trained: WT: n=22 mice; *Mecp2*-null, n=20mice. Error bars represent mean ± SE. ns, not significant, ** *P* < 0.01, *** *P* < 0.001, Two-way ANOVA test with Sidak’s Post-Hoc.

**Fig. S13.**
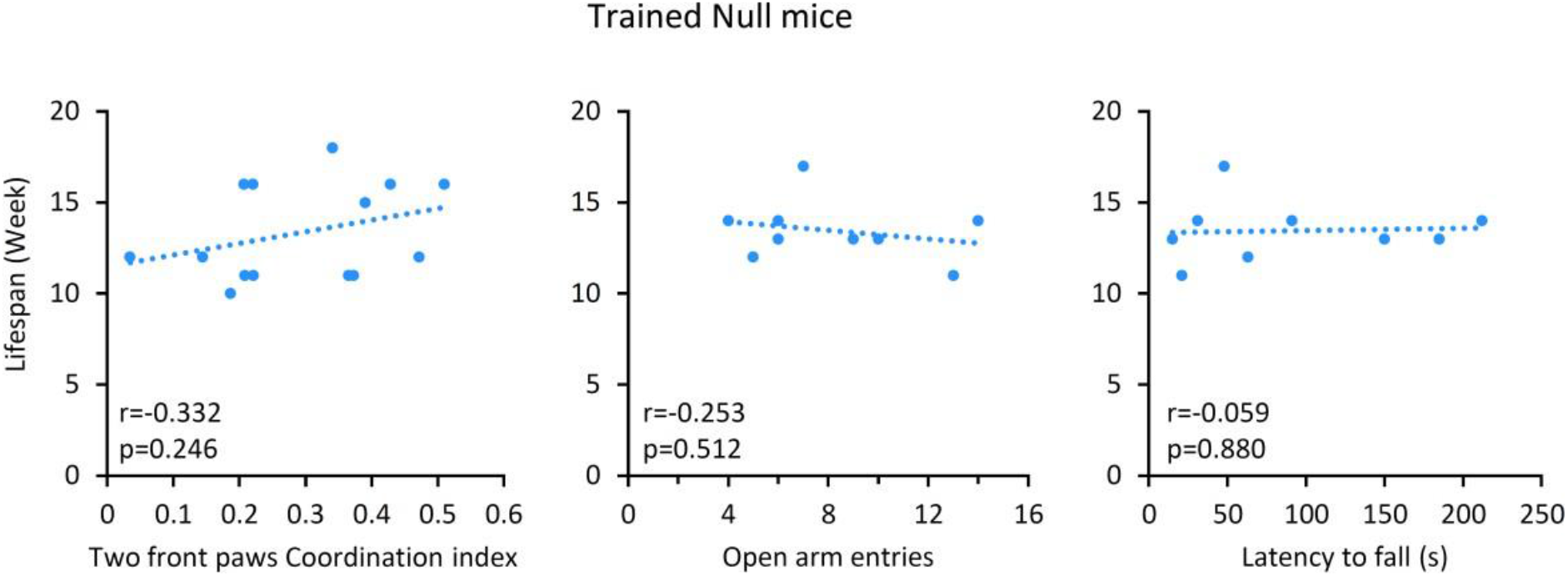
Longer lifespan correlates with motor learning, but not with reduced anxiety-like behavior or stamina on the rotarod. Although the correlation between the lifespan and improved forepaw coordination is weak (p = 0.236 and r = 0.336, Pearson bivariate correlations test), it suggests that motor learning contributes to extending the survival of *Mecp2*-null mice.

**Supplemental table 1:**
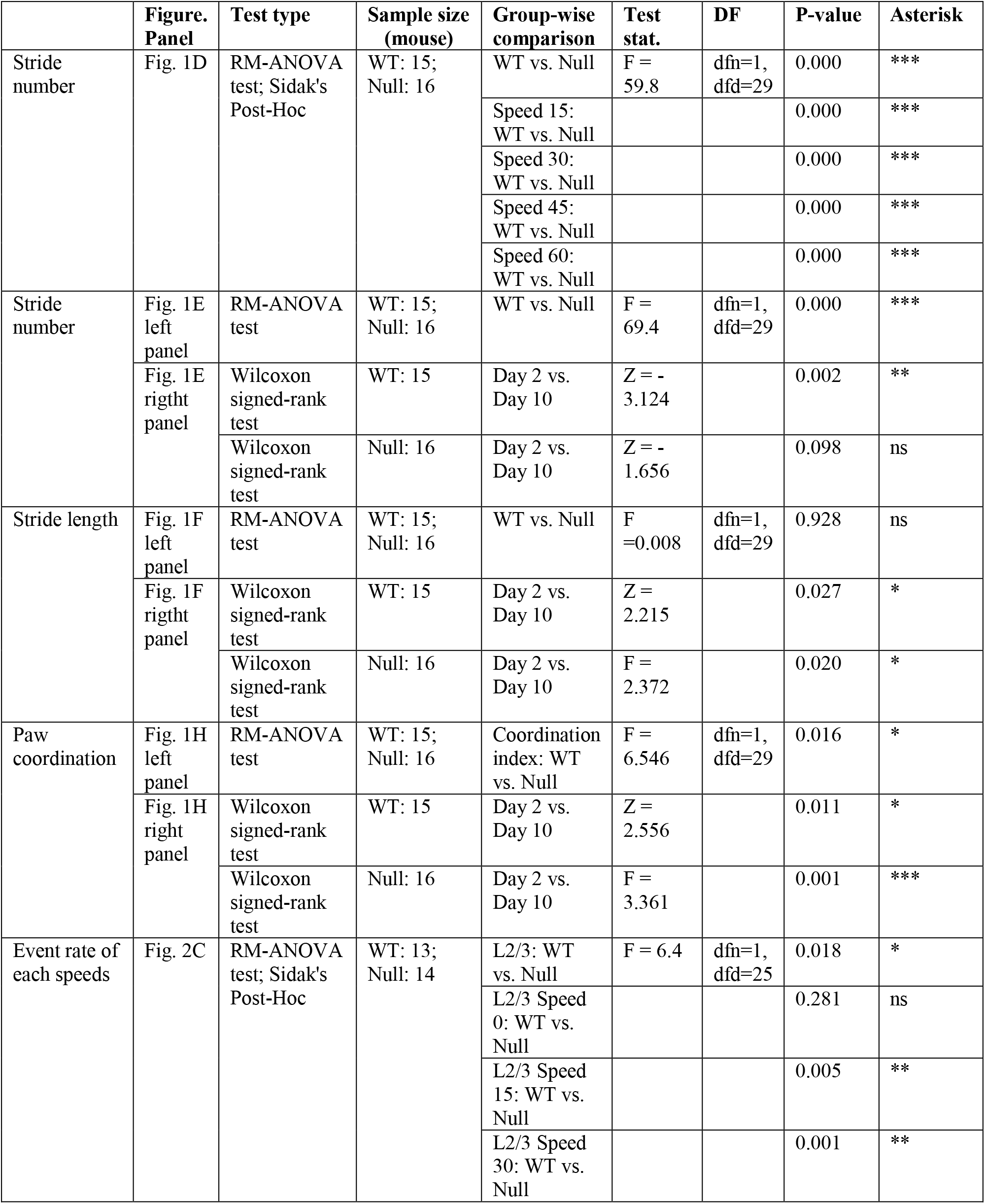
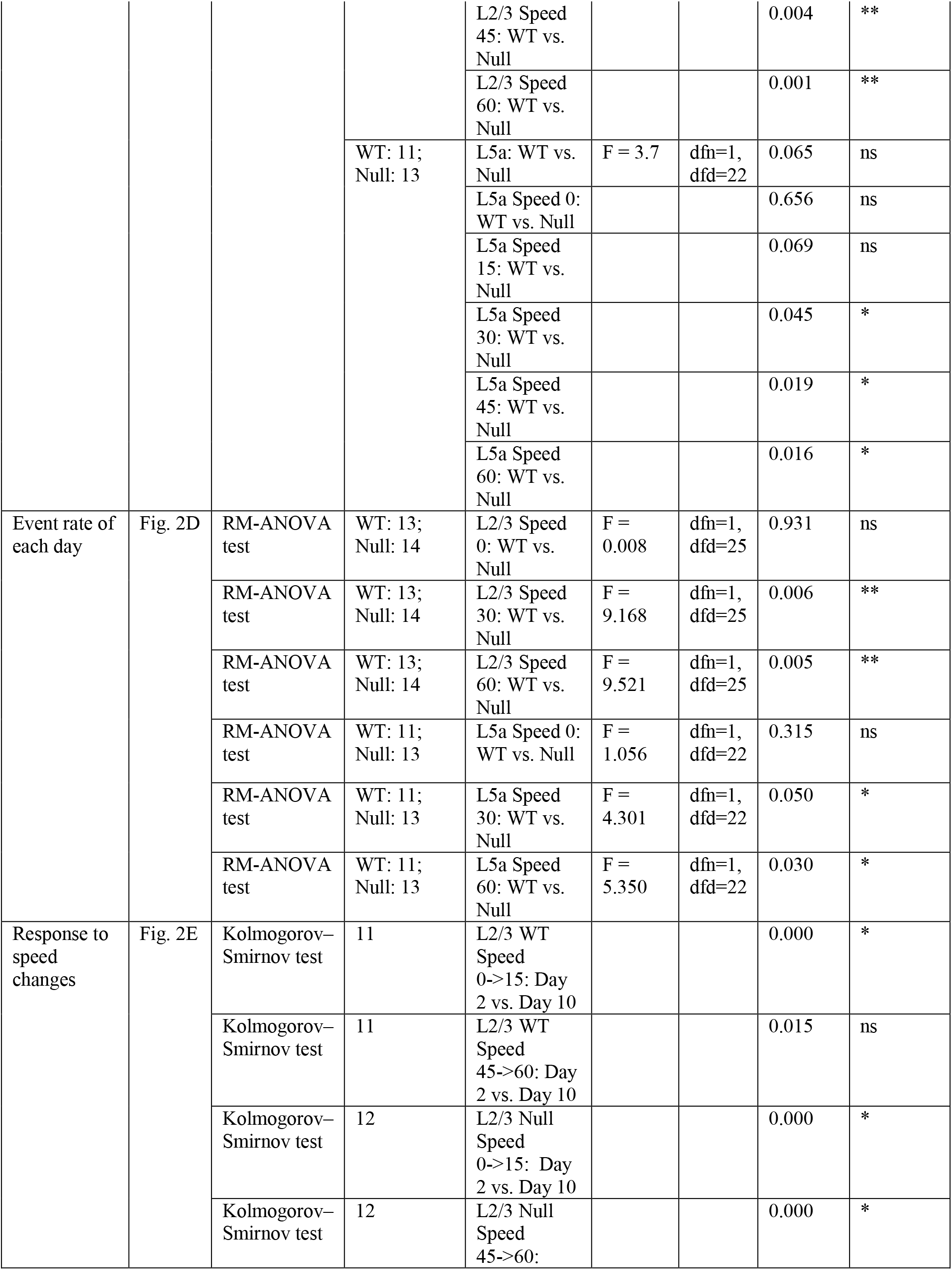
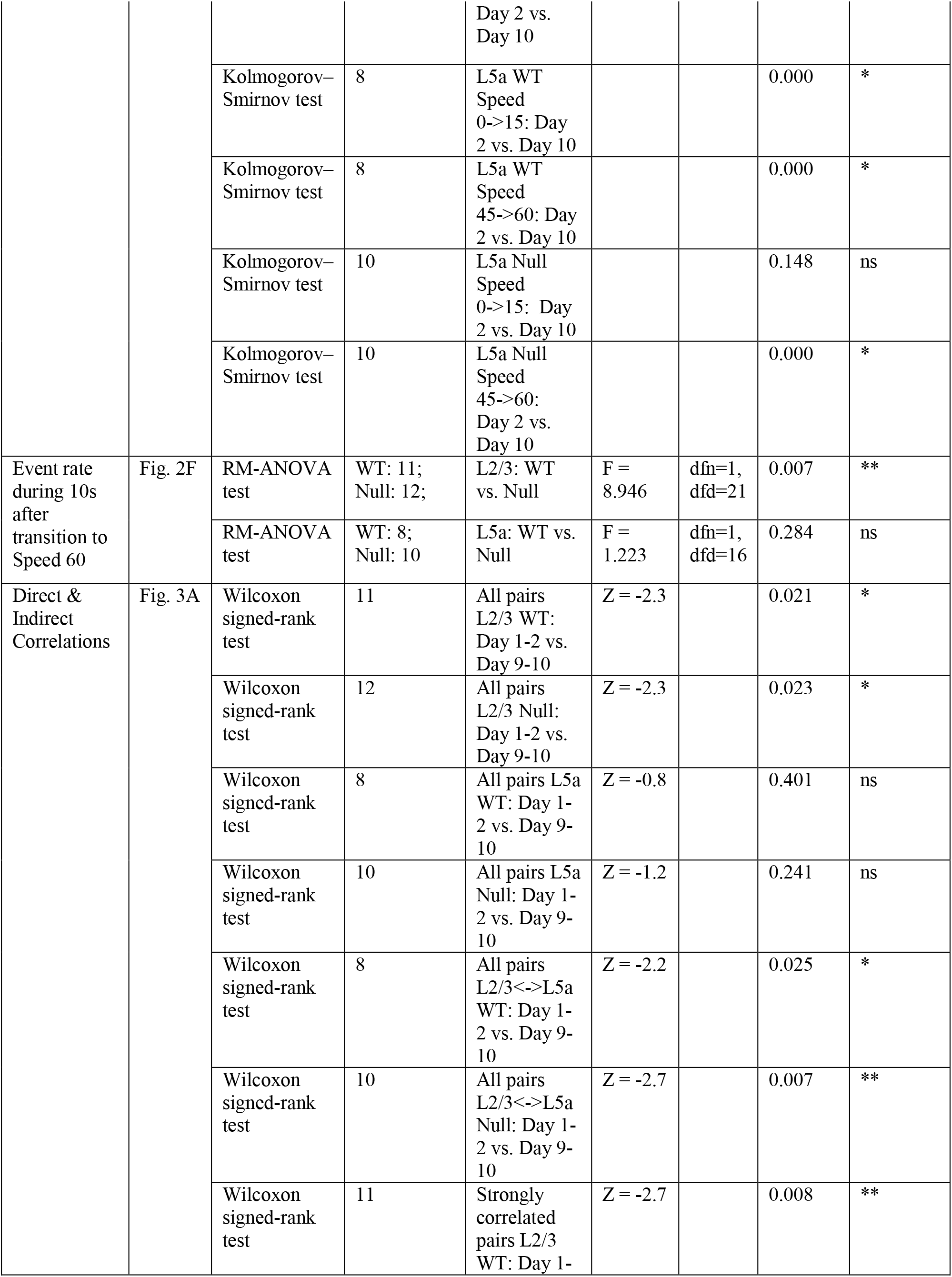
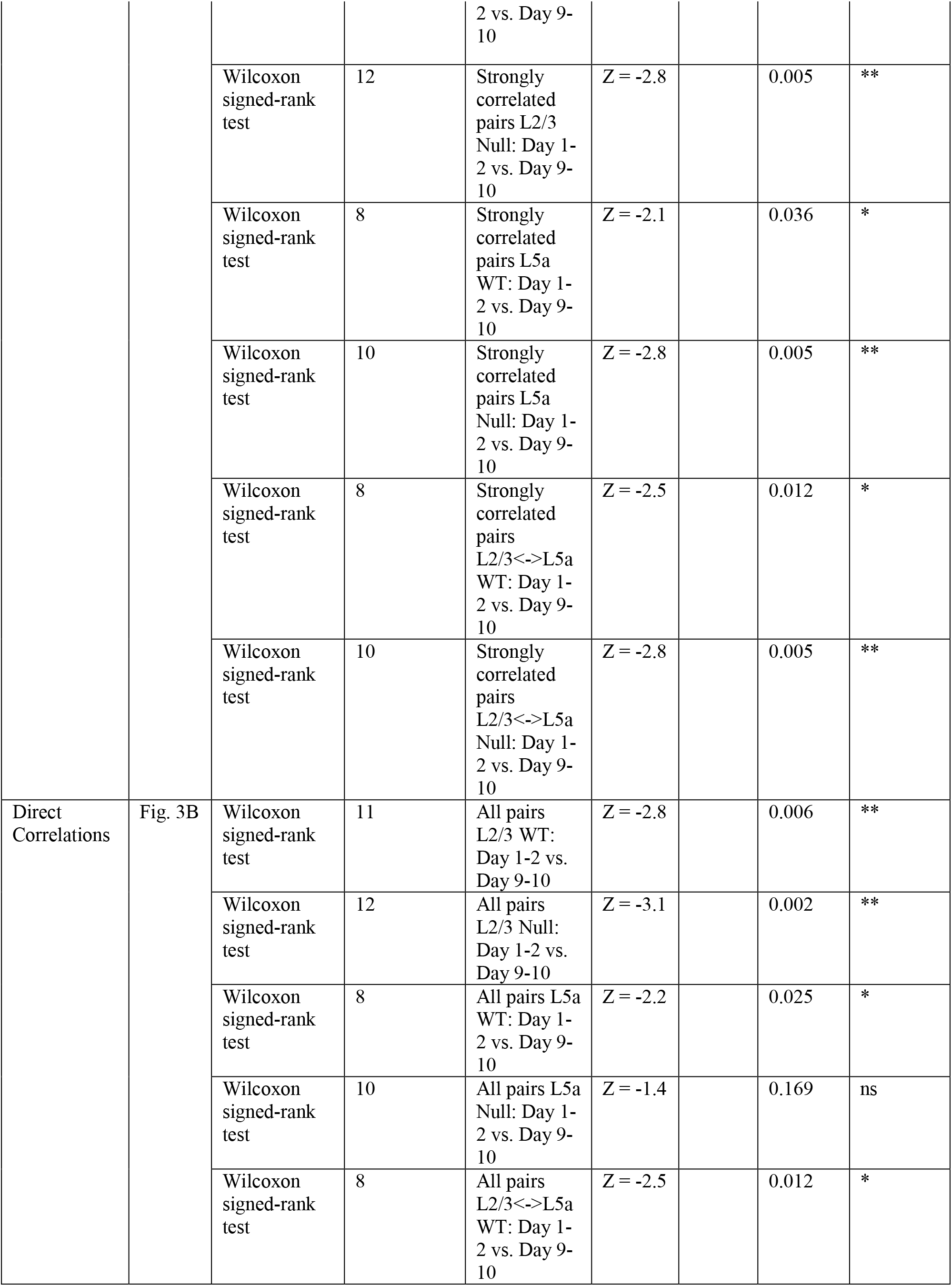
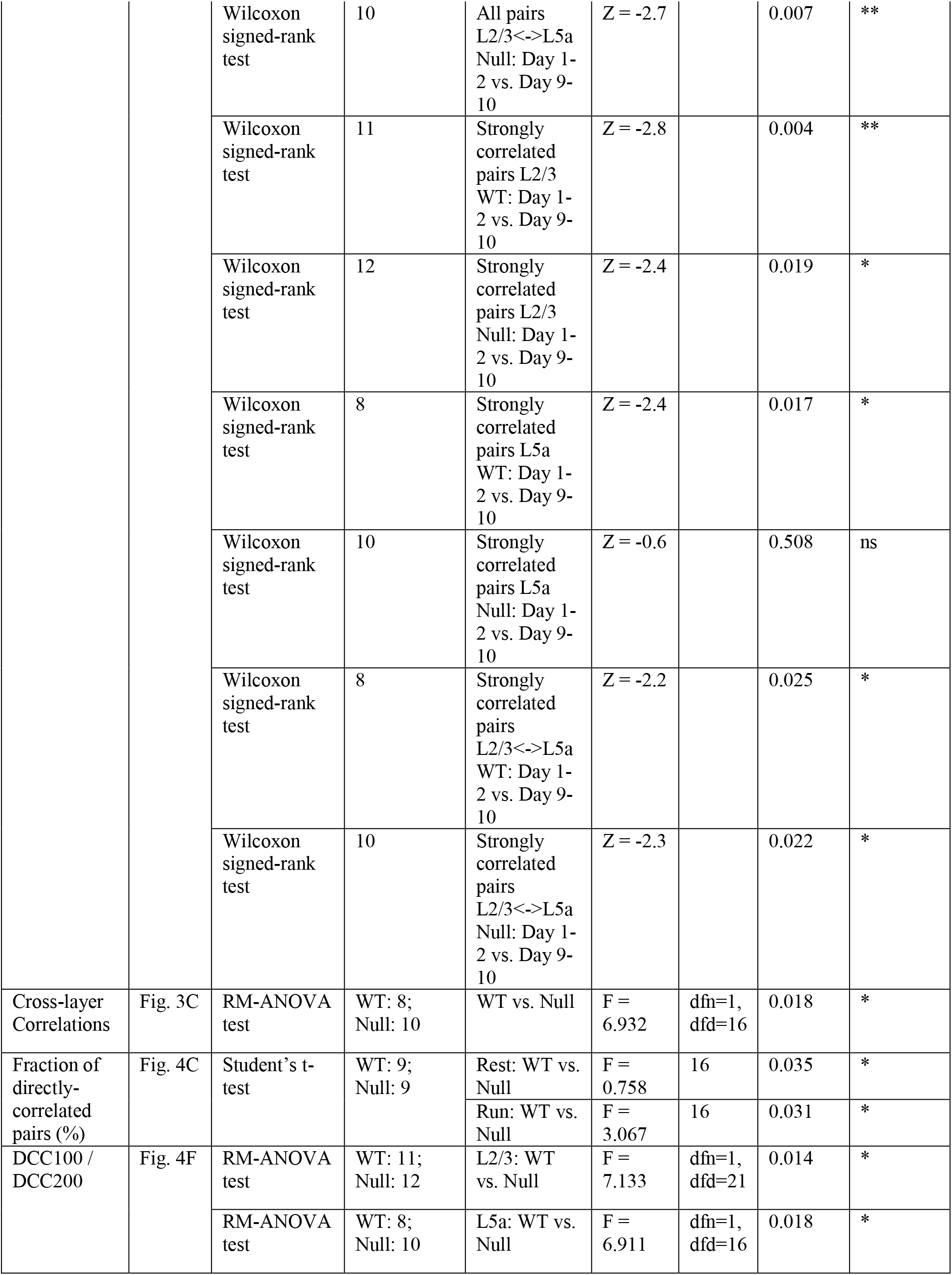
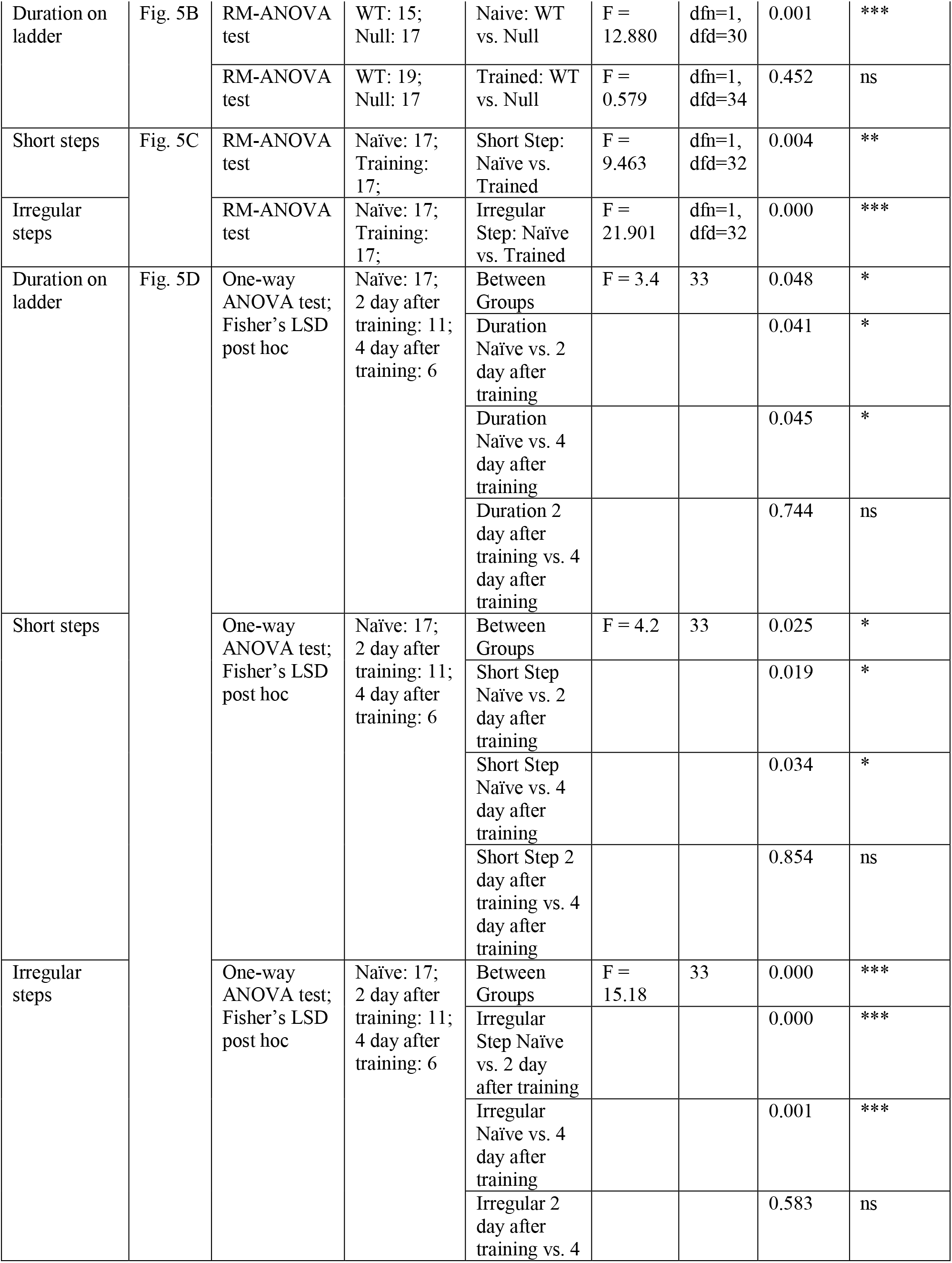
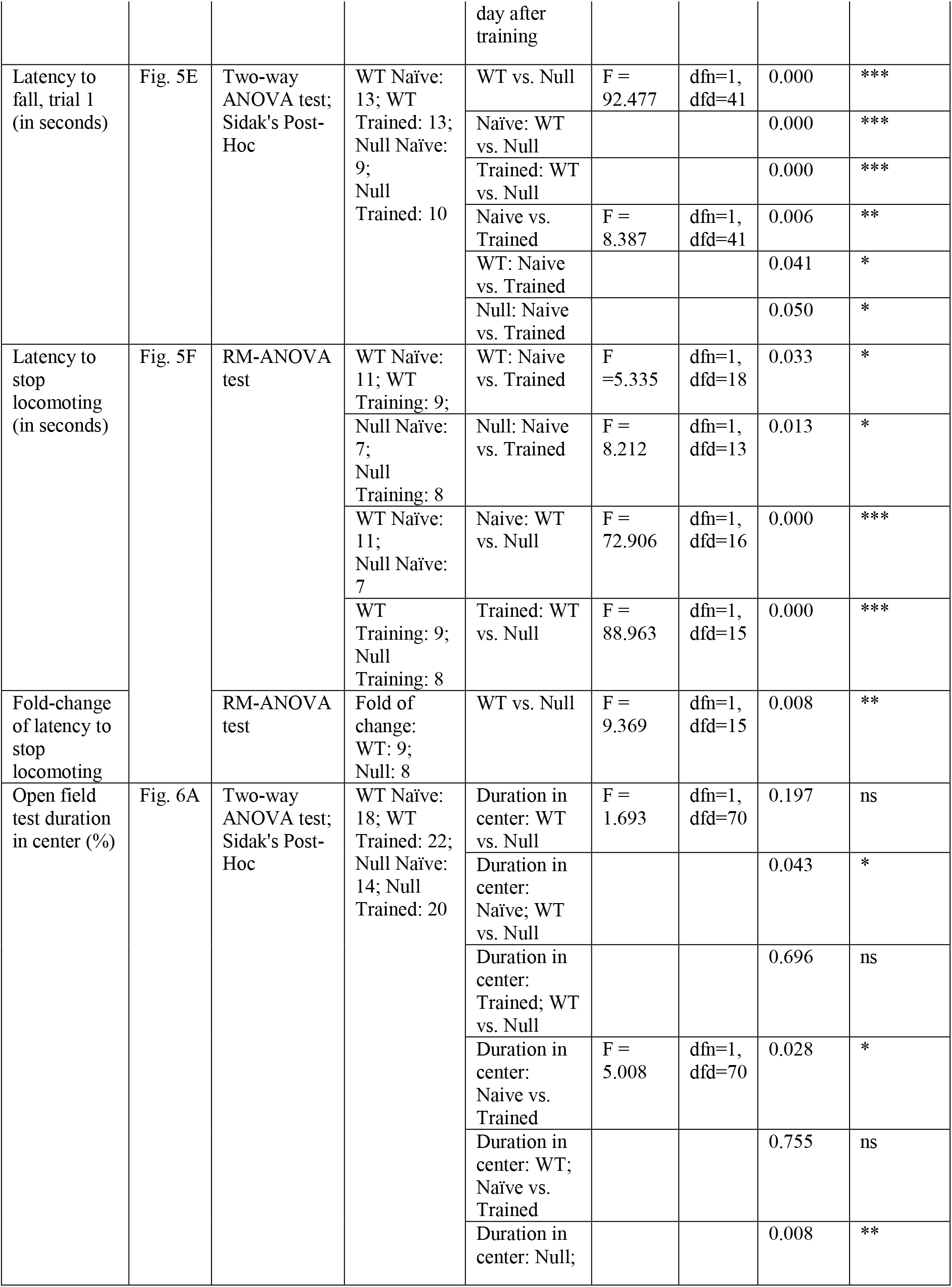
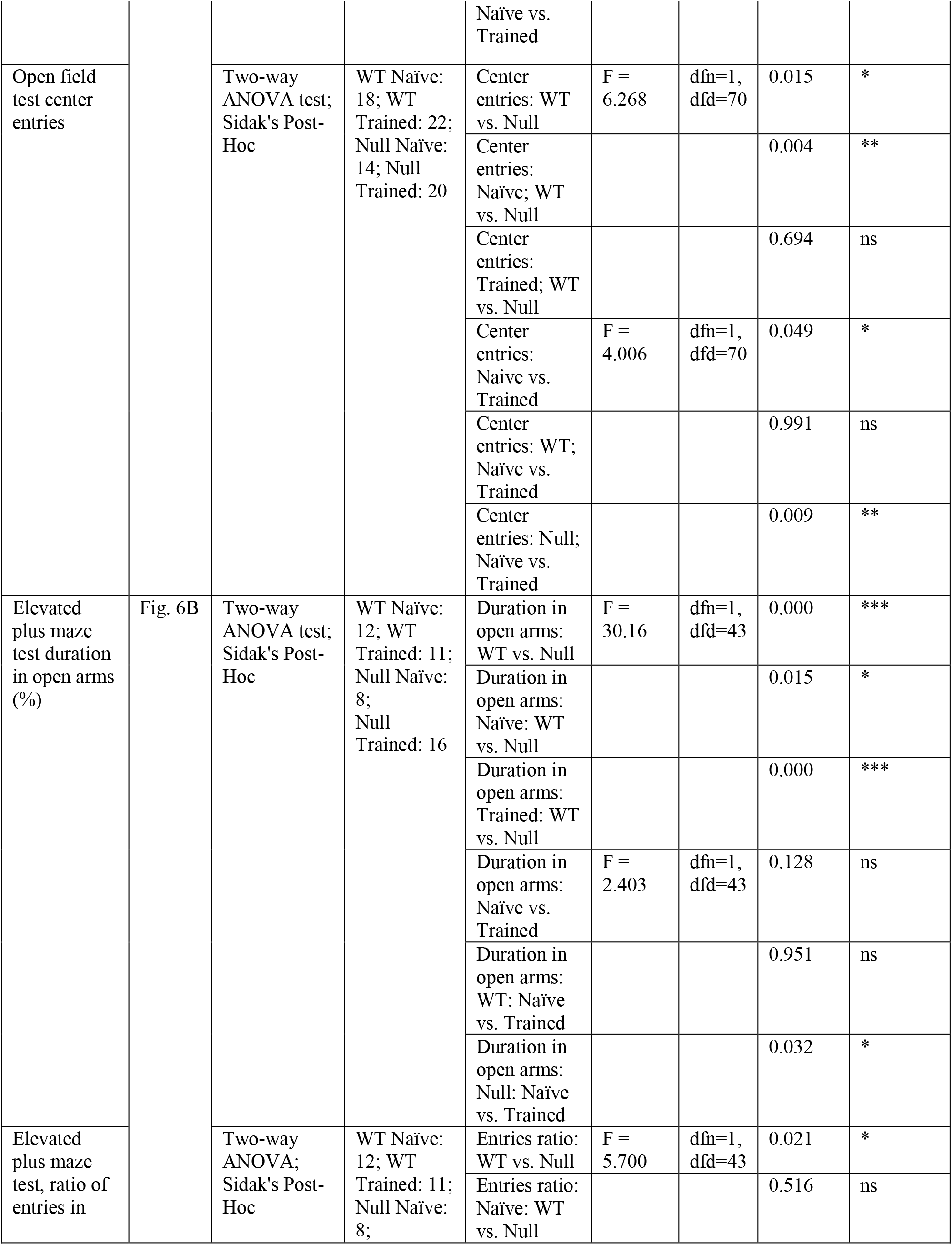
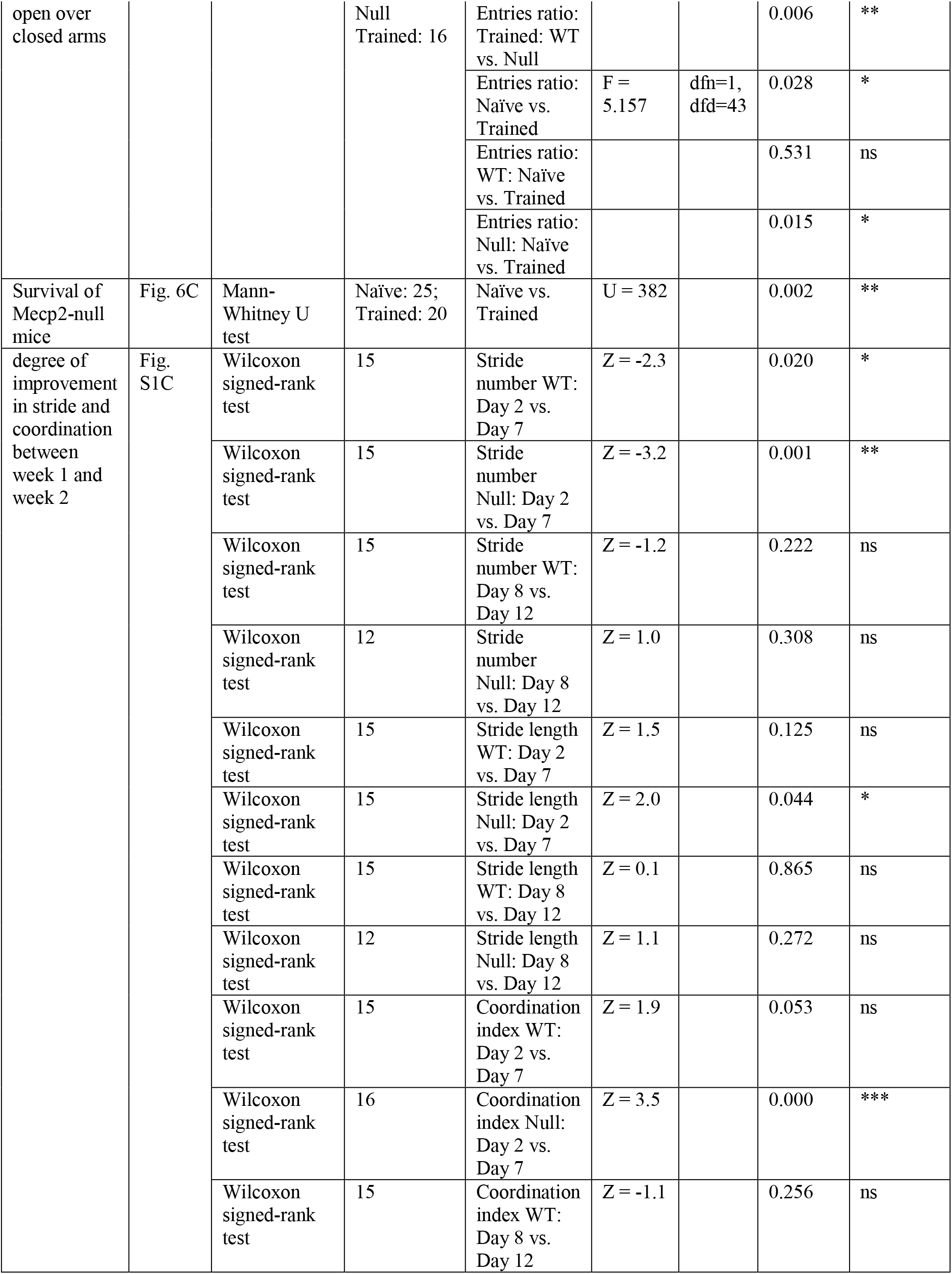
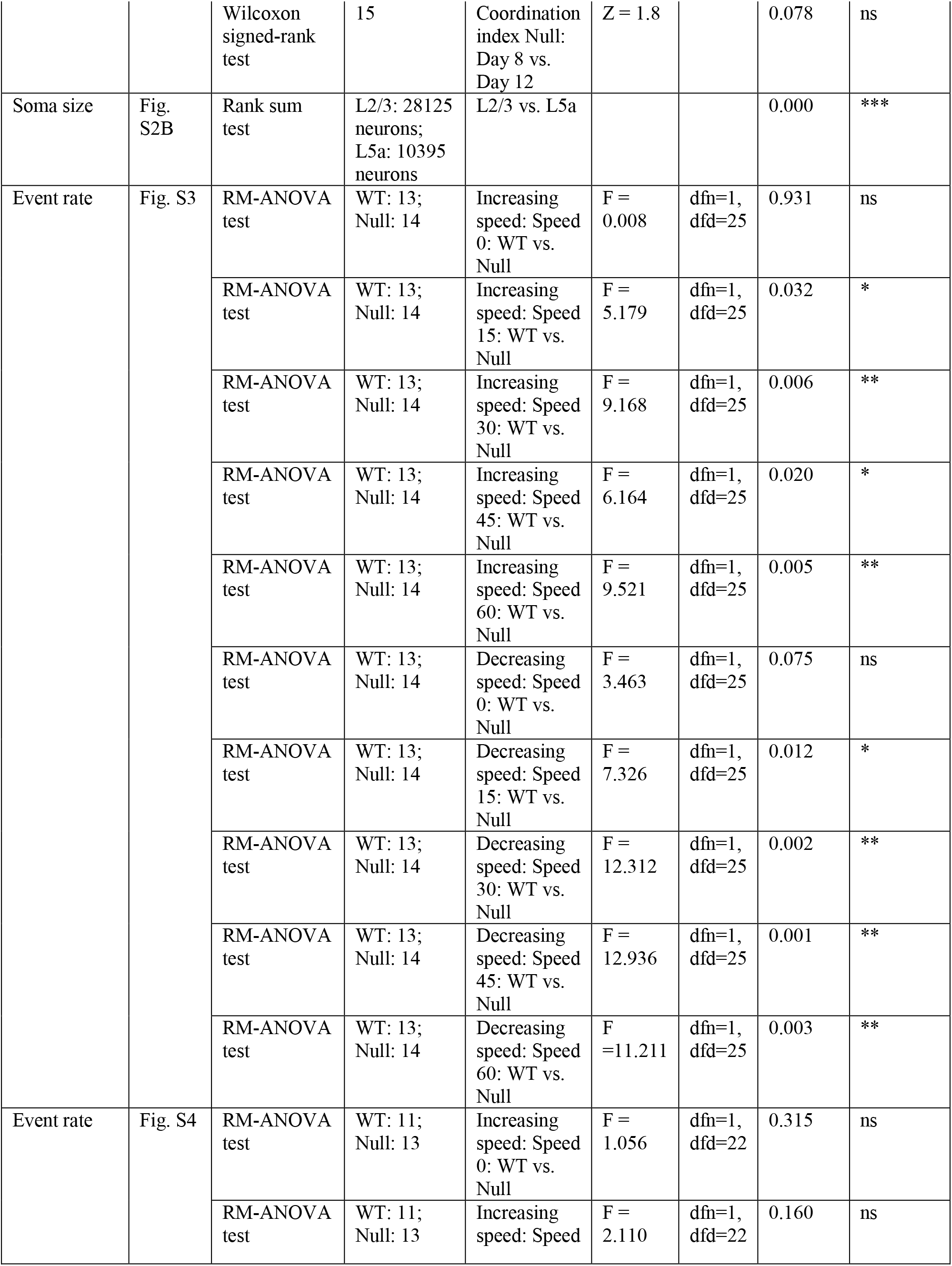
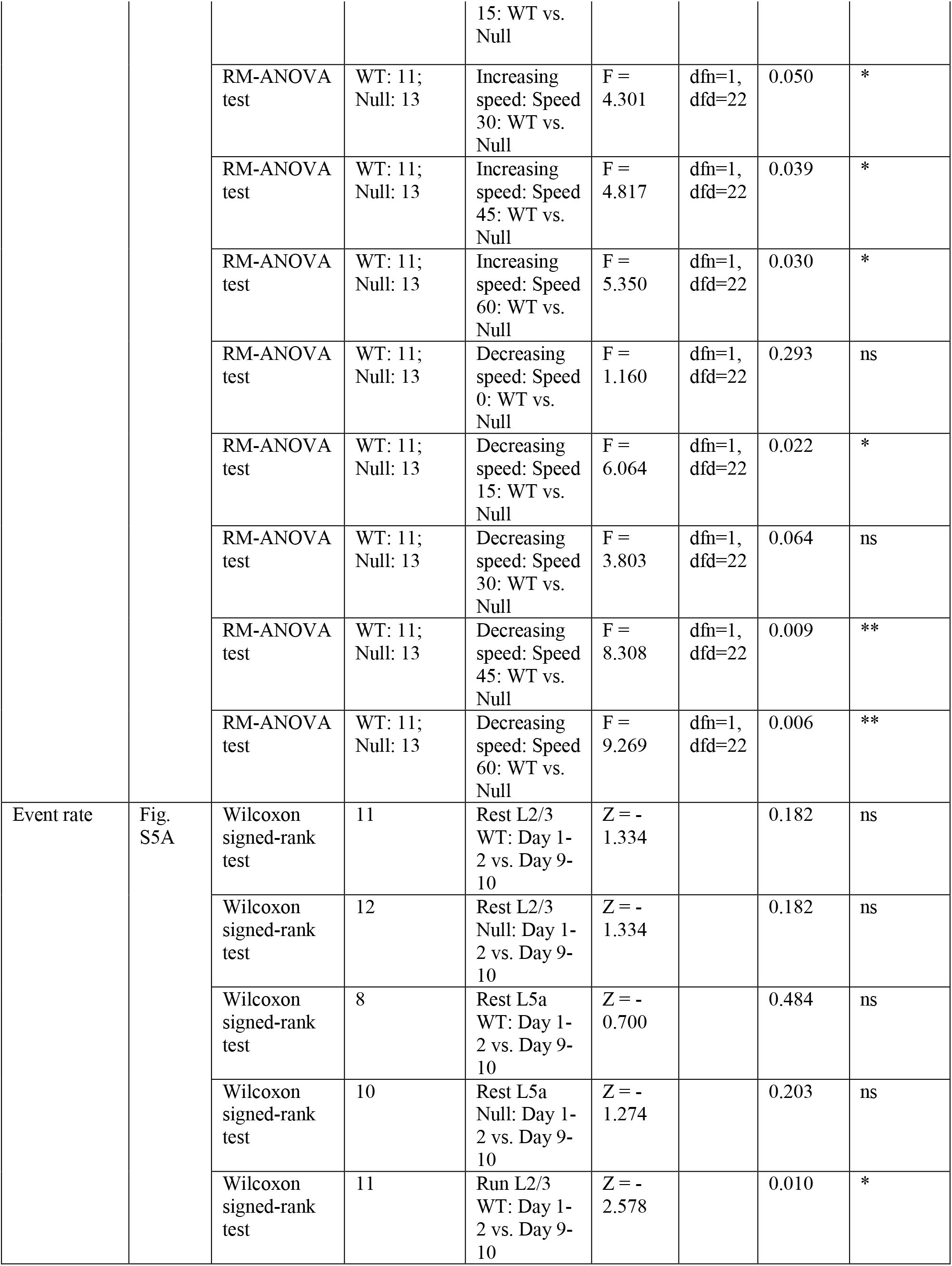
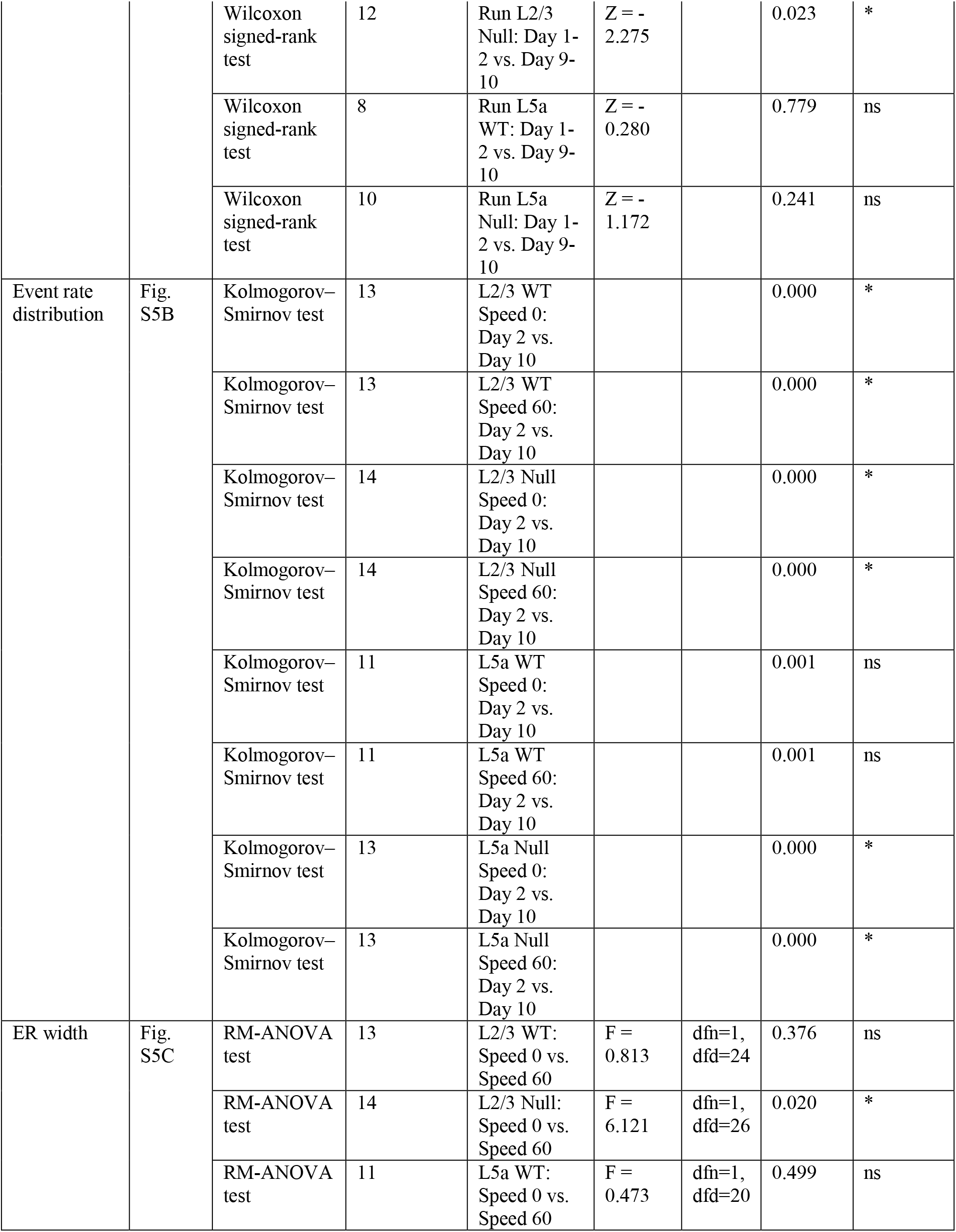
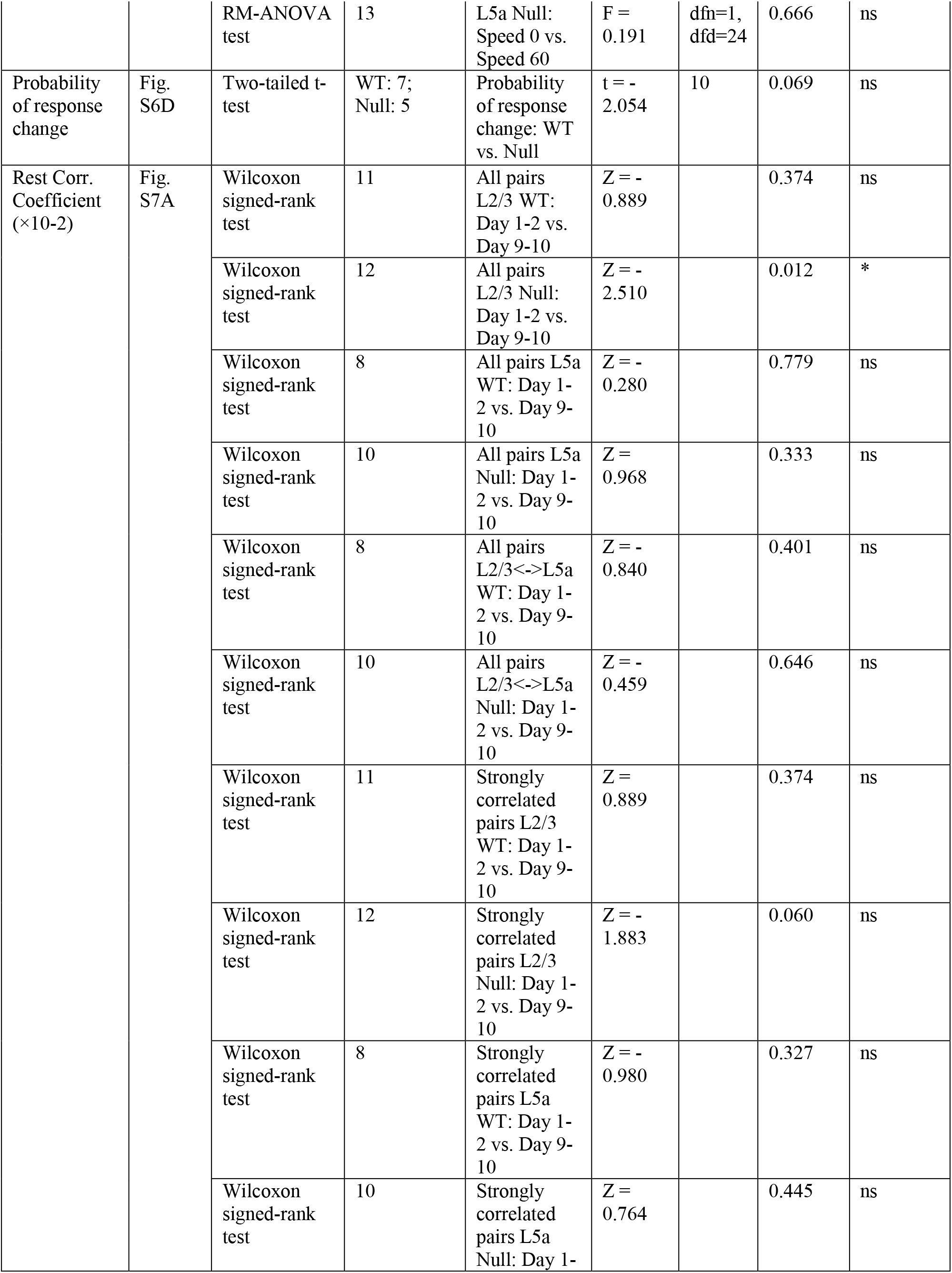
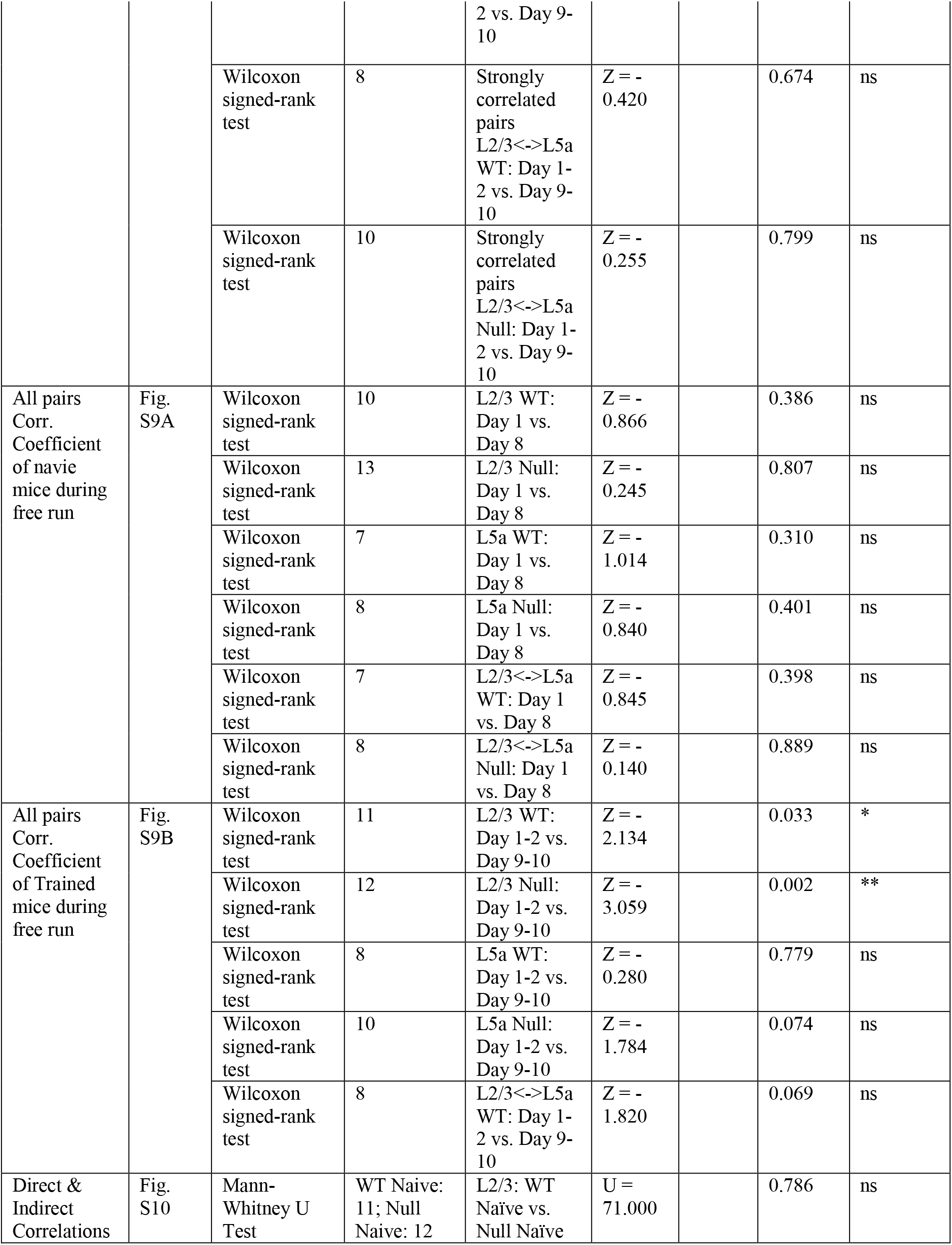
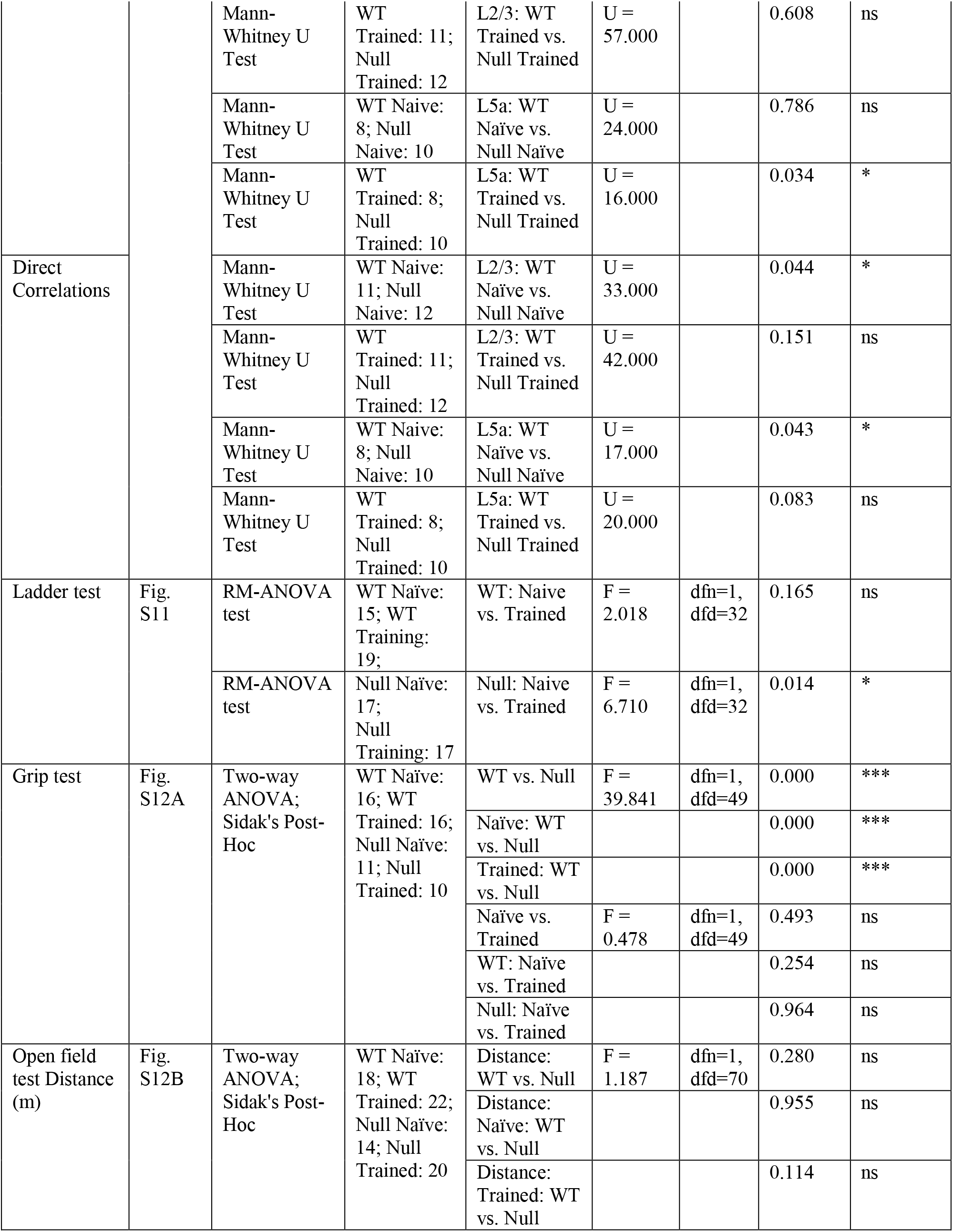
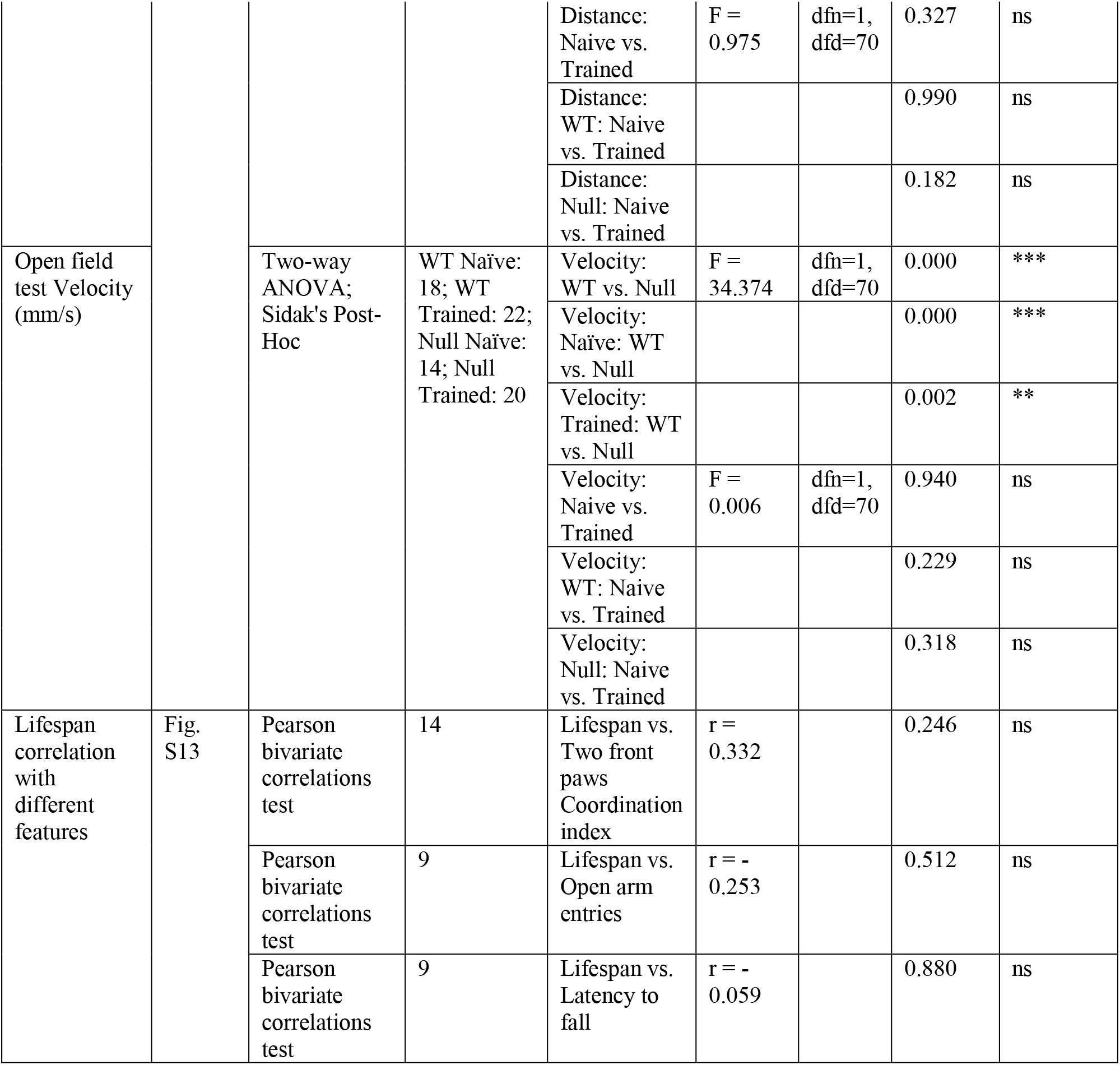
Detailed statistical results for each experiment, organized by figure panel.

